# Functional redundancy enables emergent metabolic dynamics in marine microbiomes

**DOI:** 10.1101/2025.08.12.669827

**Authors:** Sergio González-Motos, Lidia Montiel, Vanessa Balagué, Francois-Yves Bouget, Ramon Massana, Josep M. Gasol, José M. González, Pierre E. Galand, Ramiro Logares

## Abstract

Understanding how marine microbiomes will respond to ongoing global change is crucial. Functional redundancy, the capacity of different microbes to perform the same function, is considered a key mechanism underpinning the stability and resilience of the ocean microbiome. Although the extent of functional redundancy remains debated, investigating its manifestation in environmentally similar and interconnected microbial communities may provide critical insights into its role in shaping microbial community dynamics. We hypothesized that examining the long-term synchrony and rhythmicity of temperate microbial communities in such locations could provide insight into the role of functional redundancy. High functional redundancy at the community level would manifest as rhythmic and synchronous metabolic functions across sites, even in the absence of synchrony or rhythmicity at finer organizational levels, such as individual genes or taxa, thereby contributing to community resilience. Conversely, low functional redundancy would imply that synchrony and rhythmicity extend to both the contributing genes and taxa, suggesting a greater vulnerability of the community to environmental variability. To test this framework, we analyzed the long-term synchrony and rhythmicity of two marine-coastal microbiomes in the Mediterranean Sea, separated by approximately 150 km and connected by a dominant southwest current. Monthly collected metagenomes from a seven-year period were examined at the levels of metabolic functions (e.g., KEGG pathways), predicted genes (open reading frames), and taxa. We found functions, genes, and taxa exhibiting high, low, or anti-synchrony, as well as displaying rhythmic or non-rhythmic patterns. Although rhythmic behavior was observed on average across all organizational levels, consistent with the seasonal dynamics expected in temperate Mediterranean waters, average synchrony across microbiomes remained low. Focusing specifically on 45 markers of key biogeochemical functions, we revealed that several functions exhibited high synchrony and rhythmicity, in sharp contrast to the low synchrony and rhythmicity among the most abundant genes and taxa contributing to those functions. This suggests that functional redundancy and complementary dynamics at lower organizational levels, with distinct taxa contributing to key metabolic functions at different times, lead to rhythmic and synchronous dynamics at higher levels through emergent self-organization. Together, our results highlight functional redundancy and emergent self-organized dynamics as key mechanisms supporting the stability and resilience of marine microbiomes under environmental change.

## INTRODUCTION

The ocean microbiome harbors a plethora of microorganisms (bacteria, archaea, viruses, and microeukaryotes) that are fundamental for the functioning of the global biogeochemical cycles and, consequently, the biosphere (*1*, *2*). Marine microbiomes exhibit a considerable degree of spatiotemporal turnover, responding to a multitude of factors, including physicochemical gradients, nutrient availability, ocean currents, and seasonality (*3–5*). This spatiotemporal variability affects community composition, diversity, and metabolisms, making the ocean microbiome a dynamic entity. Nonetheless, in several regions and depths, this shifting nature remains broadly predictable, characterized by specific patterns of daily, seasonal, and interannual variations in community composition, which emerge as inherent properties of the system (*6*). In particular, the ocean microbiome often exhibits strong seasonal rhythmicity in temperate and polar areas, with community composition and function changing yearly in a predictable manner due to cyclical changes in environmental drivers, such as light, temperature, and nutrient availability (*7*, *8*). In several cases, the predictable dynamics of microbial communities are manifested as rhythmic patterns of microbial abundances, which can influence nutrient cycling, primary production, and other ecosystem processes (*9*). The rhythmicity of the ocean microbiome has been investigated in different locations around the globe (*10–12*). Despite significant progress, our understanding of the extent to which seasonal dynamics are synchronized across interconnected microbiomes remains limited. Gaining insights into this synchrony could illuminate whether marine microbiomes linked by currents operate as interconnected, coordinated systems or as more functionally independent entities. Comprehending synchrony could also contribute to delineating different microbial assemblages in the ocean and determining their sizes. For example, the rhythmic and synchronous presence of multiple taxa in specific locations aligns with the idea that the distribution of certain microbes follows large-scale patches across the ocean (*13*).

Seasonal synchrony refers to the coordinated and predictable growth or decline of microbes over time across assemblages in the same environmental setting (*14*). In a perfect synchrony scenario, microbial taxa are expected to grow or decline simultaneously, reaching their highest or lowest abundances at the same time of the year. In practice, differences in biotic or abiotic local and regional processes may result in lower synchrony dynamics or different rhythmic patterns. For example, pathogens or predators may stochastically increase their abundances in specific locations, affecting the rhythmicity patterns within a single community and the synchrony between communities (*15*, *16*). Furthermore, microbial synchrony and rhythmicity depend on physical processes (*17*) such as ocean connectivity mediated by currents and wind (*18*). Ocean connectivity facilitates microbial dispersal across regions, potentially enhancing synchrony and rhythmicity by increasing community similarity or integrating microbes into the same physical patch (*13*). Connectivity can also align environmental conditions across regions (e.g., characterized by temperature and nutrients), driving microbiome synchrony by homogenizing the environmental factors that shape microbial community structure (*19*). However, connectivity could also decrease microbial synchrony. For example, by introducing species or pathogens that may disrupt community dynamics (*20*). Occasionally, microbes may also respond synchronously to sporadic environmental triggers such as sudden changes in nutrient availability, temperature, algal blooms, and sediment disturbances (*21*, *22*). In this case, non-rhythmic but synchronous dynamics could be expected.

Microbial synchrony is also expected to depend on intrinsic features specific to each microbiome, such as the degree of functional redundancy. This refers to the property of different taxa sharing the same ecological functions (e.g., nutrient cycling or carbon fixation) (*23*, *24*). Functional redundancy is crucial, as it can enhance resilience by allowing the microbial community to maintain essential functions even when one or more species are removed or their population sizes are reduced (*25*). Ecologically redundant taxa do not necessarily occupy exactly the same niche since they may respond slightly differently to environmental heterogeneity. Overall, one could expect high functional redundancy to decrease the rhythmicity and synchrony of microbial communities, as fluctuations in the abundance of individual taxa are less likely to impact community functioning significantly. However, this still remains to be empirically tested.

Determining whether or not connected microbial assemblages display synchronous and rhythmic seasonal dynamics can provide insights into the factors that drive community turnover and the strength of environmental selection (*26*). Stronger selection pressure, when consistent across different locations, may potentially lead to higher synchrony (*5*). In line with this, previous studies have documented microbial synchrony across a variety of aquatic systems, particularly in freshwater environments where communities respond in unison to shared seasonal drivers. Studies in temperate rivers, circumpolar Arctic rivers, and regional lake networks have shown that both bacterial and phytoplankton communities can fluctuate synchronously over time, largely due to common environmental forcing such as temperature, hydrology, or nutrient availability (*17*, *27*, *28*). In marine environments, synchrony has been observed as tightly coordinated gene expression patterns among co-occurring populations, driven by diel cycles (*29*). However, little is known about whether such synchrony extends across larger spatial and temporal scales.

While synchrony remains largely underexplored in the ecology of marine microbes, mainly due to the lack of comparable long-term time series, there have been advances in gut microbiology (*30*, *31*). These studies have focused on gut microbiotas from individuals sharing similar environments and diets. In these cases, community changes are influenced not only by environmental factors or diet but also by variations in host genetics. Also, ocean connectivity may be several orders of magnitude greater than that observed in those individual gut microbiomes. Despite such differences, some similarities are found. For instance, at least 73% of the microbial core functionality in the ocean—essential for basic vital cellular processes—is shared with the human gut microbiome (*32*). While this does not imply a similar temporal dynamic, it may provide a common foundation for investigating how rhythmicity and synchrony arise in functionally redundant systems. Ultimately, it is essential to recognize that in both cases, variations in microbial communities are expected to arise from fundamental ecological processes (e.g., environmental selection, dispersal, and ecological drift) that broadly influence microbial dynamics. According to metacommunity theory, microbial dispersal is a key factor in promoting synchrony across communities (*14*, *33*). This idea was tested in a recent study on the gut microbiomes of wild baboons, which—despite sharing environments and diets over 13 years—reported weak synchrony among individuals. The authors suggested that microbial dispersal was too weak to overcome the primary drivers that define the distinctive features of microbiomes. Among these, functional redundancy played a key role, having more influence than dispersal (*34*). Recognising functional redundancy as a dominant feature in microbial systems aligns with marine microbiome studies showing that although the taxonomic composition of microbial communities varies tremendously across oceans, their gene composition and functional capacity are more conserved (*23*, *32*). It suggests the existence of substantial functional redundancy, although the functional level used to measure redundancy may influence the results (*24*, *35*). Altogether, the previous studies suggest that functional redundancy and limited dispersal decrease the amount of synchrony among microbiomes at least at the taxonomic (and gene) level.

Functional redundancy may shape the emergence of synchrony and rhythmicity across organizational levels within microbial communities. We hypothesize that in marine microbiomes, high functional redundancy is reflected in scenarios where metabolic functions exhibit synchronous and rhythmic dynamics over time, even though the specific genes or taxa contributing to these functions, representing finer levels of biological organization, do not. Conversely, low functional redundancy would be indicated when both functions and the associated genes or taxa demonstrate synchrony and rhythmicity over time. This raises the question of the extent to which the dynamics of a given function, and thus its synchrony and rhythmicity, can be explained by the dynamics of its associated genes (ORFs) or taxa. In the ocean, many metabolic processes are often dominated by highly prevalent taxa (*36*), so they strongly influence the dynamics of the function to which they contribute. However, in some cases, rhythmicity may emerge at the functional level despite the absence of rhythmic patterns in the individual contributing genes (ORFs), indicating that rhythmic functional dynamics can arise from the combined activity of non-rhythmic genes. For example, vitamin B_12_ is an essential factor regulating the growth of photosynthetic planktonic communities (*37–41*) and an indispensable molecule for many auxotrophs of this vitamin (*42*, *43*). In a recent study conducted in a 7-year monthly metagenomic time series in the Mediterranean Sea, genes (ORFs) annotated as *cobF*, *cobG,* and *cobA-btuR* exhibited non-rhythmic dynamics yet contributed to rhythmic function-level B_12_ dynamics, thus demonstrating functional redundancy at the gene level (*44*).

Here, we investigate the role of functional redundancy in shaping the long-term structure of marine microbial communities by analyzing patterns of synchrony and rhythmicity in two neighboring marine microbial assemblages from the northwestern Mediterranean Sea. These sites, the Blanes Bay Microbial Observatory (BBMO) and the Banyuls Bay Microbial Observatory (SOLA), are approximately 150 km apart and are broadly connected by a dominant southwestward current (Figure 1A). Both microbiomes experience similar environmental heterogeneity and strong seasonality, with day length and temperature significantly influencing microbial community dynamics. To explore functional redundancy, we analyze synchrony and rhythmicity at different organizational levels, including metabolic functions, genes (ORFs), and taxa. We ask whether neighboring, interconnected marine microbial assemblages exhibit synchronous and rhythmic dynamics, whether these patterns vary across levels of biological organization or trait types, and whether some of these dynamics reflect emergent self-organization arising from functional redundancy.

**Figure 1.**
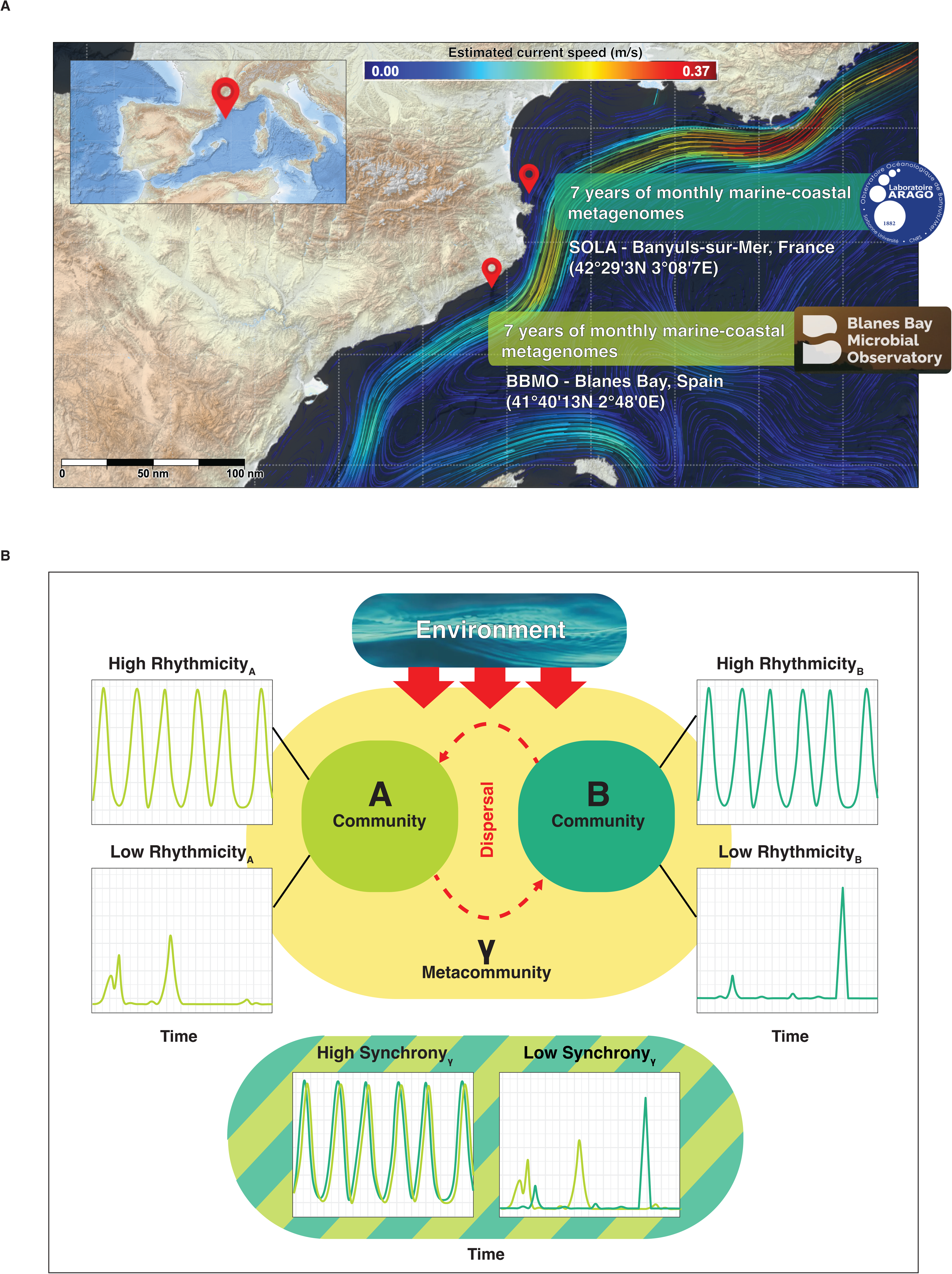
Theoretical framework and geographical overview on how synchrony may emerge in connected ocean microbiomes, indicating rhythmic or non-rhythmic dynamic features (functions, genes, or OTUs). Panel. **A.** Map illustrating the coastal monthly sampling locations BBMO (41°40’13N 2°48’0E) and SOLA (42°29’3N 3°08’7E) in the northwestern Mediterranean Sea. The red-blue color palette shows the estimated current’s speed in m/s (upper color bar) of the Liguro-Provençal-Catalan Current flowing from north to southwest and potentially connecting both microbiomes. **Panel B.** Two microbial communities are depicted: Community A (e.g., BBMO) and Community B (e.g., SOLA). Both communities are part of a broader metacommunity (γ), and are potentially connected via dispersal (exchange of organisms or genetic material, represented by the curved arrows). Environmental drivers (e.g., temperature, nutrients) influencing each site are indicated with straight arrows labelled Environment. The four smaller plots show theoretical examples of microbial dynamics over time (x-axis: Time, e.g., monthly sampling over several years), with the y-axis representing the relative abundance of microbial features. These plots illustrate high or low rhythmicity (seasonal recurrence) in each community, and how these patterns may result in high or low synchrony at the metacommunity level (γ).

## RESULTS

### Long-term microbial dynamics at two neighboring time series

We analyzed 93,303 functions from eggNOG, 16,624 from Pfam, 4,872 from COG, 8,455 from KEGG, and 429 from CAZy databases, as depicted in **Table 1**. Overall, the average synchrony was in the lower range (mean = 0.23) (**Figure 2A**), and functions from all databases showed similar synchrony distributions. More precisely, most functions (61.08%) displayed low synchrony, while 16.25% exhibited negative synchrony. Only a small proportion of the functions (3.90%) exhibited high synchrony, while 18.75% displayed medium synchrony. All these values represent the mean across databases. Functions were also classified according to metabolic pathways to which they belonged, as organized by eggNOG, KEGG, and CAZy databases, but no apparent differences were observed between the considered ranges of synchrony (**Figure S1**).

**Figure 2.**
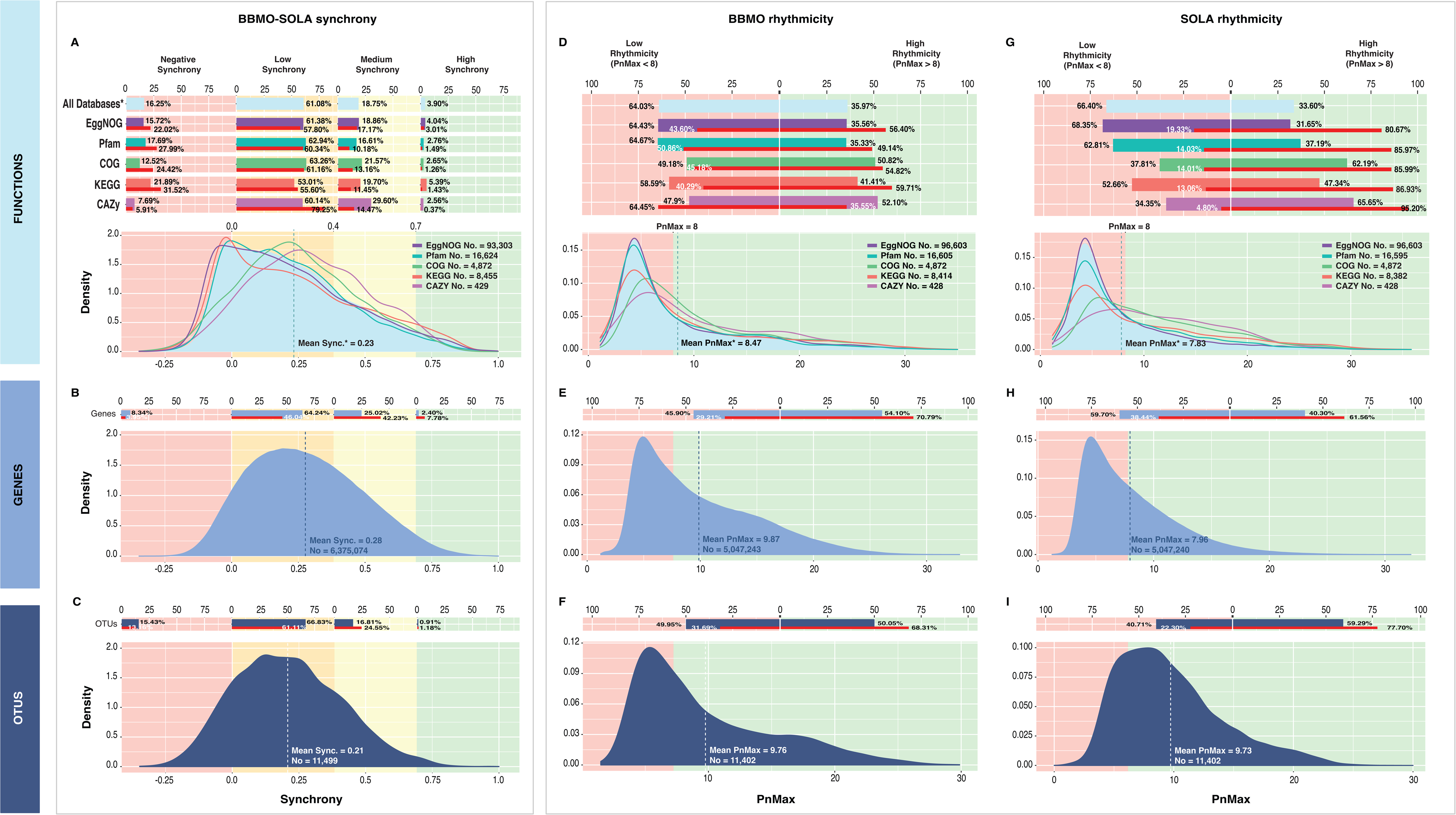
BBMO and SOLA displayed average rhythmic patterns (PnMax > 8) at the three considered levels: metabolic functions, genes (ORFs), and OTUs. However, on average, they showed low synchrony across all levels (Sy ∼ 0.25). Panel. **A.** Density plot of functions annotated with eggNOG, Pfam, COG, KEGG, and CAZy with a total number of 123,683 functions analysed (Across all databases), categorized by synchrony degree: high (Sy ≥ 0.7), moderate (0.4 ≤ Sy < 0.7), and low (Sy < 0.4). Percentages for each database and for the summatory (All Databases) within the estimated criterion are depicted above in horizontal bars. Red horizontal bars indicate the percentage they represent of the total abundance for all panels. **Panel B.** Density plot of 6,375,074 genes, each with at least 30% occurrence throughout the dataset, according to the established synchrony degree criterion and with percentages above. **Panel C**. Density plot of 11,499 OTUs (mTags) with at least 30% of occurrence throughout the dataset according to the synchrony degree established boundaries, and with percentages above. **Panel D.** Density plot of BBMO functions with a total number of 122,228 functions analysed according to the rhythmicity (PnMax) threshold and with percentages above. **Panel E.** Density plot of 5,047,243 BBMO genes with at least 30% of occurrence throughout the dataset according to the rhythmicity (PnMax) threshold and with percentages above. **Panel F.** Density plot of 11,402 BBMO OTUs with at least 30% of occurrence throughout the dataset according to the rhythmicity (PnMax) threshold and with percentages above. **Panel G.** Density plot of SOLA functions with a total number of 122,196 functions analysed according to the rhythmicity (PnMax) threshold and with percentages above. Note that the rhythmic functions in SOLA represent the vast majority of the abundance with respect to BBMO rhythmic functions. **Panel H.** Density plot of 5,047,240 SOLA genes with at least 30% of occurrence throughout the dataset according to the rhythmicity (PnMax) ranges and with percentages above. **Panel I.** Density plot of 11,402 SOLA OTUs with at least 30% of occurrence throughout the dataset according to the rhythmicity (PnMax) threshold and with percentages above.

**Table 1.**
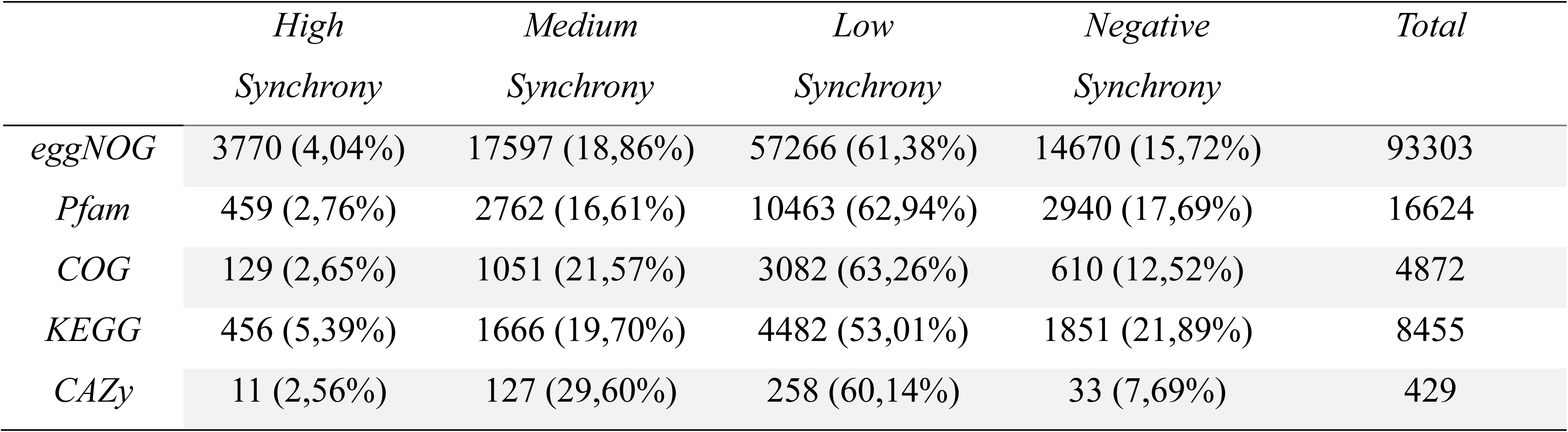
Synchrony of functions specified by eggNOG, Pfam, COG, KEGG and CAZy databases in the BBMO-SOLA time series.

|  | *High*<br>*Synchrony* | *Medium*<br>*Synchrony* | *Low*<br>*Synchrony* | *Negative*<br>*Synchrony* | *Total* |
| --- | --- | --- | --- | --- | --- |
| *eggNOG* | 3770 (4,04%) | 17597 (18,86%) | 57266 (61,38%) | 14670 (15,72%) | 93303 |
| *Pfam* | 459 (2,76%) | 2762 (16,61%) | 10463 (62,94%) | 2940 (17,69%) | 16624 |
| *COG* | 129 (2,65%) | 1051 (21,57%) | 3082 (63,26%) | 610 (12,52%) | 4872 |
| *KEGG* | 456 (5,39%) | 1666 (19,70%) | 4482 (53,01%) | 1851 (21,89%) | 8455 |
| *CAZy* | 11 (2,56%) | 127 (29,60%) | 258 (60,14%) | 33 (7,69%) | 429 |

At the gene level (6,375,074 ORFs), there was a lower proportion of medium and high synchronous genes (25.02% and 2.40%, respectively) than genes featuring negative and low synchrony (8.34% and 64.24%, respectively). Yet, medium and high synchronous genes represented a larger fraction of the total abundance, reaching values of 42.23% and 7.78%, respectively. This means that the more abundant genes tend to exhibit higher synchrony. (**Figure 2B**; see horizontal red bars). Similar patterns were observed for OTUs. Overall, out of 11,499 OTUs, those with low synchrony prevailed, representing 66.83% of the total, with an average synchrony of 0.21 (**Figure 2C**). Only a small portion, comprising 0.91% of the OTUs, exhibited high synchrony. Similarly to genes (ORFs), the most abundant OTUs displayed high synchrony.

As a general pattern, functions, genes, and OTUs tended to be rhythmic in both microbiomes, particularly when considering their relative abundances. However, we also found functions, genes, and OTUs showing rhythmic and arrhythmic patterns in both microbiomes. At the functional level in BBMO, for instance, most functions were not rhythmic when considering their number alone, with 64.43% in eggNOG, 64.67% in Pfam, and 58.59% in KEGG functions (**Table 2**). In turn, 50.82% of COG functions and 52.10% of CAZy functions exhibited high rhythmicity (**Table 2**). Nevertheless, highly rhythmic functions accounted for a proportionally greater share of the total abundance: 56.40% in eggNOG, 49.14% in Pfam, 54.82% in COG, and 59.71% in KEGG. The only exception was CAZy, where highly rhythmic functions represented 35.55% of the total abundance. On average, for functions from all databases, BBMO functions manifested rhythmic patterns as the PNmax value of 8.46 was higher than the threshold established in the literature to define rhythmicity (PNmax ≥ 8) (*20*, *44*) (**Figure 2D**).

**Table 2.** Rhythmicity of BBMO functions corresponding to eggNOG, Pfam, COG, KEGG and CAZy databases.

|  | *Low Rhythmicity*<br>*(PnMax < 8)* | *High Rhythmicity*<br>*(PnMax > 8)* | *Total* |
| --- | --- | --- | --- |
| *eggNOG* | 62246 (64,43%) | 34357 (35,57%) | 96603 |
| *Pfam* | 10739 (64,67%) | 5866 (35,33%) | 16605 |
| *COG* | 2396 (49,18%) | 2476 (50,82%) | 4872 |
| *KEGG* | 4930 (58,59%) | 3484 (41,41%) | 8414 |
| *CAZy* | 205 (47,90%) | 223 (52,10%) | 428 |

In SOLA, we also observed that, in bulk, most functions were not rhythmic when considering their number alone, yet highly rhythmic functions accounted for a disproportionately large share of the total abundance. Specifically, 68.35% of eggNOG, 62.81% of Pfam, and 52.66% of KEGG functions exhibited low rhythmicity (**Table 3**). On the contrary, 62.19% of COG and 65.65% of CAZy functions displayed high rhythmicity (**Table 3**). However, highly rhythmic functions accounted for a larger share of the total abundance: 80.67% in eggNOG, 85.97% in Pfam, 85.99% in COG, 86.03% in KEGG, and 95.20% in CAZy. Overall, in SOLA, highly rhythmic functions represented a remarkably higher proportion of the total abundance than in the BBMO (56.40% eggNOG, 49.14% Pfam, 54.82% COG, 59.71% KEGG, and 35.55% CAZy). On average, for all databases, SOLA functions displayed an average PNmax of 7.83, which falls slightly below the threshold established in the literature to determine rhythmicity (PNmax ≥ 8) (*20*, *44*) (**Figure 2G**). Nevertheless, when all databases were considered collectively, rhythmic functions accounted for approximately 86.77% of the total abundance, underscoring the pronounced rhythmicity of SOLA functions.

**Table 3.**
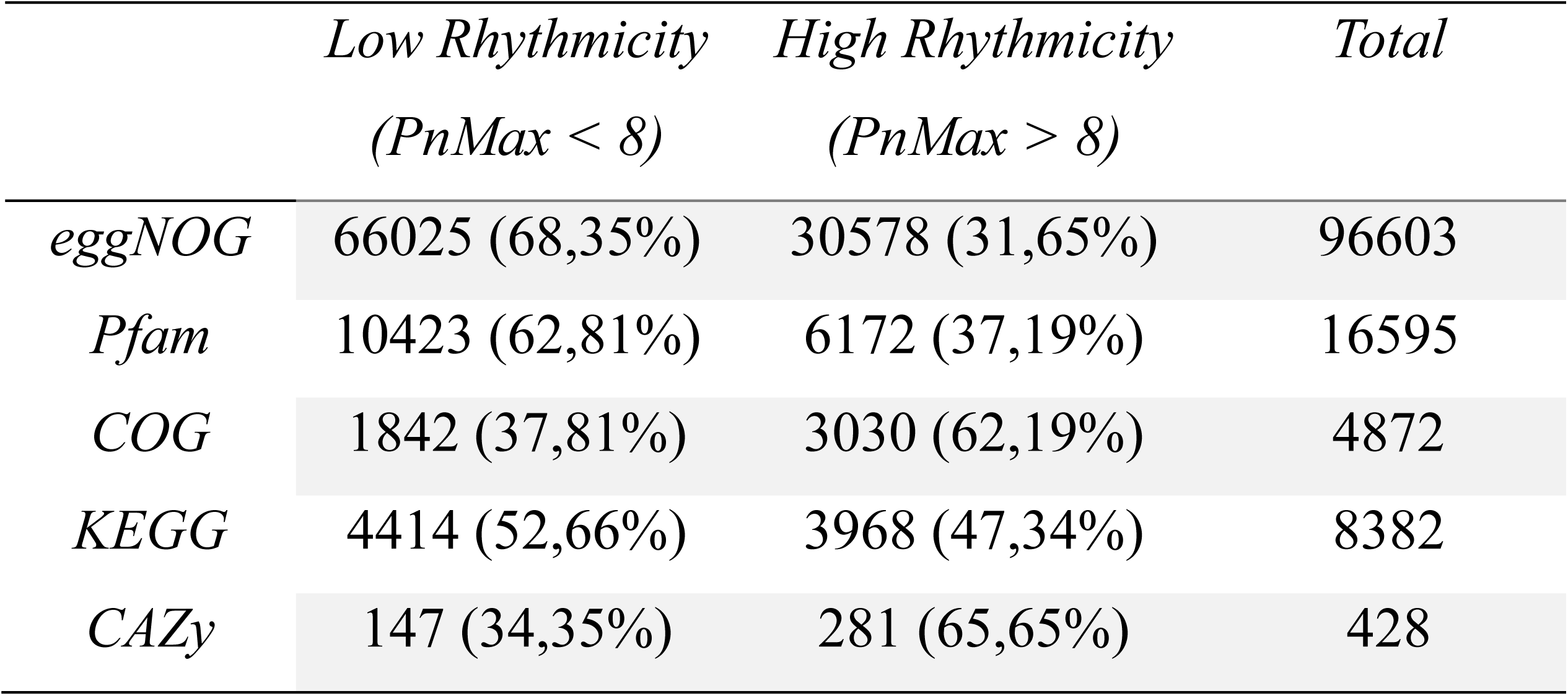
Rhythmicity of SOLA functions corresponding to eggNOG, Pfam, COG, KEGG and CAZy databases.

At the gene (ORF) level, from a total of 5,047,243 genes analyzed in BBMO, 45.90% displayed low rhythmicity, accounting for 29.21% of the total abundance. In contrast, 54.10% showed high rhythmicity, representing 70.79% of the total abundance. The mean PNmax of all genes was 9.87, depicting high rhythmicity (**Figure 2E**). In the case of SOLA, 59.70% displayed low rhythmicity, accounting for 38.44% of the total abundance. In turn, 40.30% showed high rhythmicity, representing 61.56% of the total abundance. The mean PNmax of all genes was 7.95 (**Figure 2H**). Despite being slightly below the established threshold of 8, SOLA genes (ORFs) were considered rhythmic since genes with a PNmax above 8 represented the major part of the total abundance (61.56%). Overall, the proportion of highly rhythmic genes was higher in BBMO (54.10%) than in SOLA (40.30%). Likewise, the representation of rhythmic genes based on their abundance contribution was higher in BBMO (70.79%) than in SOLA (61.56%).

Regarding taxa, from a total of 11,402 OTUs in BBMO, 49.95% displayed low rhythmicity, accounting for 31.69% of the total abundance. In turn, 50.05% of the OTUs exhibited substantial rhythmicity (PNmax ≥ 8), which accounted for 68.31% of the overall abundance. The mean PNmax across all OTUs was 9.76 (**Figure 2F**). In SOLA, from a total of 11,402 OTUs, 40.71% displayed low rhythmicity, accounting for 22.30% of the total abundance. In contrast, 59.29% displayed a substantial degree of rhythmicity, representing 77.70% of the overall abundance. The average PNmax across all OTUs was 9.73 (**Figure 2I**). The abundance contribution of rhythmic OTUs was proportionally similar in both BBMO and SOLA microbiomes.

### Dynamics of biogeochemical key functions

We investigated the temporal dynamics of a subset of 45 biogeochemical functions crucial in the carbon, nitrogen, phosphorous, and sulfur metabolisms, as well as for H_2_ oxidation and other iron-related processes (**Figure 3)**. We observed functions exhibiting contrasting dynamics in both microbiomes, that is, synchronous and asynchronous, as well as rhythmic and non-rhythmic (**Figure 3A**). Functions in the carbon and nitrogen metabolism, such as *coxL, rbcS, chlG, amoA, amoC, amoB,* and *nirK* showed high synchrony and rhythmicity **(Figure 3A)**. Synchrony and rhythmicity were correlated across all functions for all BBMO genes (Pearson correlation coefficient π = 0.5129, p-value <0.001, R^2^= 0.2631) and for all SOLA genes (π = 0.5143, p-value <0.001, R2= 0.2646). Therefore, synchronous functions were also commonly rhythmic, indicating comparable environmental selection in both sites. This aligns with the fact that most samples from BBMO and SOLA displayed a similar environmental heterogeneity (**Figure S2**).

**Figure 3.**
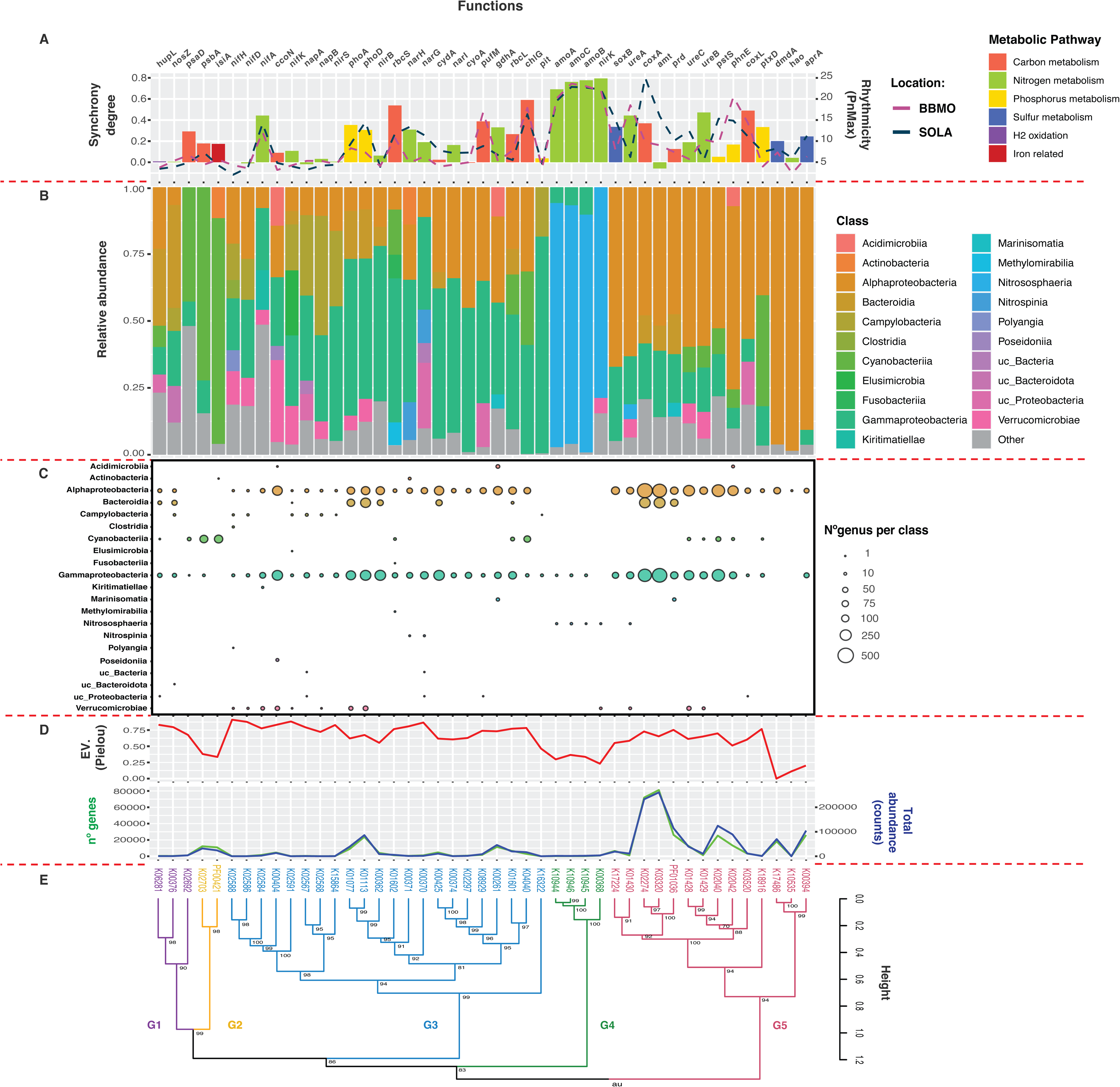
Synchrony, rhythmicity, and taxonomic distribution of biogeochemically key functions in BBMO and SOLA. Panel. **A.** Barplot showing the synchrony degree and rhythmicity of each selected function specified in coloured dotted lines, all arranged according to the corresponding metabolic pathway (indicated with colours). **Panel B.** Taxonomic relative abundance (class level) of each selected function depicted in colours. **Panel C.** Number of genera per taxonomic class for each function. Note that colours are the same as those used in B. **Panel D.** Top: Pielou’s evenness index for genes annotated in each function. Bottom: Number of genes (Y axis-left) in green and their total abundance as counts (Y axis-right) in blue for each function. **Panel E.** Euclidean hierarchical clustering of the functions according to their taxonomic composition. We show five clusters grouped at a height of 1 (indicated with colours). A p-value of each cluster is indicated as an approximately unbiased (AU) value.

However, some specific functions, such as the *amt* gene (ammonium uptake), had a rhythmic pattern in SOLA (maximum peak = 15.88) and, to a lesser extent, in BBMO (maximum peak = 9.00), but did not exhibit synchronicity (synchrony degree = -0.05). Regardless of the number of genes (ORFs) and their total abundance, functions tended to be more rhythmic and synchronous the more uneven they were in terms of ORF content, as indicated by Pielou’s evenness index. This was the case for *amoA, amoC, amoB* (ammonia oxidation), and *nirK* (denitrification) (**Figure 3D**). The previous functions were dominated almost entirely by class Nitrosophaeria (**Figure 3B**), including very few genera (**Figure 3C**). We also observed rhythmic and synchronous functions with less taxonomic dominance. This was illustrated by cases such as *rbcS* (RuBisCO, small chain), with five classes, including Alphaproteobacteria (22 genera), Cyanobacteria (9 genera), Fusobacteria (1 genus), Gammaproteobacteria (51 genera), and Methylomirabilia (2 genera) featuring a Pielou’s evenness index of 0.76. Another case was *chlG* (chlorophyll and bacteriochlorophyll *a* synthesis), driven by three different classes, including Alphaproteobacteria (84 genera), Cyanobacteria (84 genera), and Gammaproteobacteria (17 genera) with a Pielou’s evenness index of 0.78.

The selected functions were clustered into five groups based on their taxonomic composition, offering a comparative framework to facilitate interpretation rather than to represent a strict biological classification (**Figure 3E**). Group 1 (G1) was mainly composed of functions contributed by Bacteroidia and Actinobacteria. One example is the hydrogen oxidation function *hupL*, which also received notable contributions from Cyanobacteriia, Gammaproteobacteria, and unclassified Proteobacteria. Then, for *nosZ*, a function within the nitrogen metabolism, there were contributions from Campylobacteria, Gammaproteobacteria, and unclassified Bacteroidota. The most differentiated function within this group was *psaD* (photosystem I, subunit II), mediated by Cyanobacteria (especially the *Synechococcus* genus), and Gammaproteobacteria. Both synchrony and rhythmicity were low in the aforementioned functions. The second group (G2; **Figure 3E**) included functions from the carbon metabolism, like *psbA* (photosystem II P680 reaction center D1 protein), driven mostly by *Synechococcus*, and iron-related, such as *isiA*, driven by Actinobacteria and *Synechococcus*. Both functions exhibited low synchrony and rhythmicity in their respective time series. The third group (G3; **Figure 3E**) considered functions related to nitrogen metabolism, including *nifH*, *nifD*, *nifA*, *nifK*, *napA*, *napB*, *nirS*, *nirB*, *narH*, *narG*, *narI,* and *gdhA*. In addition, functions from the carbon metabolism, such as *ccoN*, *pufM*, *rbcL*, *rbcS, chlG*, *cydA,* and *cyoA*, as well as functions from the phosphorus metabolism, like *phoA*, *phoD,* and *pit*. Contributors to G3 were mainly from Gammaproteobacteria and Alphaproteobacteria. G3 displayed heterogeneous synchrony and rhythmicity. Key functions in nitrification, such as *amoA*, *amoB*, *amoC*, and denitrification, like *nirK* were clustered together in a fourth group (G4; **Figure 3E**) dominated by Archaea of the class Nitrososphaeria, in particular by the genus *Nitrosopumilus* and *Nitrosopelagicus*. G4 was the most rhythmic and synchronous group of functions, in addition to being the most uneven in terms of taxonomy. Lastly, the fifth group (G5; **Figure 3E**) included functions from the sulfur metabolism like *soxB*, *dmdA,* and *aprA*; functions from the nitrogen metabolism like *ureA*, *ureB*, *ureC*, *amt*, and *hao*; functions from the carbon metabolism like *coxA*, *coxL,* and *prd,* and functions from the phosphorus metabolism such as *pstS*, *phnE*, and *ptxD* with contributions mainly from Alphaproteobacteria, including Pelagibacter, particularly for sulfur-related functions like *aprA*. Synchrony and rhythmicity varied within the group, demonstrating considerable variability among functions.

The taxonomic composition of the studied functions in BBMO and SOLA over seven years was largely coherent with that observed in the global ocean (from the *Tara* Oceans cruise). The Pearson correlation (PPC) between BBMO-SOLA and data from the global ocean displayed a mean coefficient for all functions of 0.62 (**Figure S3A**). However, only 12 functions were statistically significant, including *psbA*, *phoA*, *phoD*, *nirB*, *narH*, *cydA*, *amoB*, *soxB*, *coxA*, *amt*, *prd,* and *aprA*. All of them show correlations close to 1 (**Figure S3A**). As expected, due to the large geographic coverage, the global ocean exhibited higher taxonomic diversity contributing to the functions than BBMO-SOLA, including additional classes (**Figure S3B**). Among them, we found Bacteriovoracia within *nifA*; Brocadiae within *hao*; Dehalococcoidia within *psbA*; Desulfobulbia within *nifD*; Desulfovibriona within *nifD*; Gracilibacteria within *nifK*; Phycisphaerae within *nifH* and *nifK* and Planktomycetia within *nirB*. However, certain taxonomical groups remained clearly dominant both in BBMO-SOLA and the global ocean, such as Nitrososphaeria in *amoA*, *amoB*, *amoC* and *nirk*; *Gammaproteobacteria* in *phoA*, *phoD*, *nirB*, *narH*, and *cydA*; or *Alphaproteobacteria* in *soxB*, *coxA*, *amt*, *prd*, and *aprA*. The number of genera per class detected in *Tara* was consistent with that observed in BBMO-SOLA (**Figure S3C**). Thus, functions such as *coxA* exhibited an elevated number of genera per class in both BBMO-SOLA and *Tara*, with 476 Alphaproteobacteria, 255 Bacteroidia, and 348 Gammaproteobacteria in *Tara*, compared to 423, 227, and 305, respectively, in BBMO-SOLA. Another example of a highly diverse function is *amt,* which exhibited 405 Alphaproteobacteria, 240 Bacteroidia, and 449 Gammaproteobacteria genera in *Tara*, compared to 366, 214, and 400 genera, respectively, in BBMO-SOLA. It is remarkable that both *coxA* and *amt* are among the most abundant functions and have the highest number of associated genes in both BBMO–SOLA and global ocean datasets (**Figure S3D**). Finally, as observed in BBMO-SOLA, the Pielou index was independent of the number of genes and total abundance, and all three indices depicted similar values to those observed in BBMO and SOLA. For example, consistent low evenness patterns were observed in *amoA* (0.27), *amoB* (0.40), and *amoC* (0.40), which closely aligned with those found in BBMO-SOLA (0.29, 0.33, and 0.36, respectively). Similar values for *amoA*, *amoB,* and *amoC* might be explained by the syntenic arrangement of these genes in the most abundant archaea (*45*). Overall, this indicates that the findings from BBMO and SOLA largely mirror broader oceanic patterns.

Our results suggest that for most key biogeochemical functions, higher degrees of synchrony and rhythmicity are present in taxonomically uneven functions, where a few taxa dominate. Environmental drivers generate recurring seasonal patterns, often led by the most abundant microbes (*7*, *8*). These, given their abundance, exert a higher influence on the dynamics of specific community functions. However, we found evidence of synchrony and rhythmicity in several less-dominated functions. This suggests that self-organization processes within species-level dynamics (lower organizational level) drive the emergence of synchrony and rhythmicity in broader metabolic functions (higher organizational level).

### Insights into functional redundancy

To better understand how functional-level patterns emerge over time, we compared the synchrony and rhythmicity of each function with those of its most abundant contributing genes (ORFs). We found 21 functions with higher synchrony than the mean synchrony of their most abundant genes or ORFs (accounting for 70% of the function abundance), in BBMO and SOLA. These included *nirK*, *narG*, *narH*, *narI*, *aprA*, *phoA*, *ureB*, *rbcL*, *rbcS*, *coxA*, *napB*, *nifA*, *psaD*, *coxL*, *chlG*, *pufM*, *amoA*, *amoB*, *amoC*, *soxB*, and *ptxD*. Similarly, we detected 21 functions with higher rhythmicity than the mean rhythmicity of their most abundant genes or ORFs in both time series. These included *nirK*, *narG*, *narH*, *aprA*, *phoA*, *phoD*, *ureC*, *rbcS*, *pstS*, *phnE*, *coxA*, *nifA*, *amt*, *coxL*, *chlG*, *pufM*, *amoA*, *amoB*, *amoC*, *soxB,* and *prd* (**Figure 4**). In total, 14 functions shared both features, that is, higher synchrony and rhythmicity than the corresponding mean of their most abundant genes (**Figure 4**). This was the case for the functions within the nitrogen metabolism *nirK, narG, narH, narI, napB, nifA*, *amoA, amoB,* and *amoC*, as well as those within the carbon metabolism *rbcS, coxA*, *coxL, chlG,* and *pufM* (**Figure 4**, red circles). In specific functions, we found very high synchrony and rhythmicity, even though the most abundant genes or ORFs contributing to these functions exhibited low mean synchrony and rhythmicity (**Figure 4**). This is exemplified by *nirK,* which had a synchrony of 0.79, while the mean synchrony of its most abundant genes was 0.29. Similarly*, nirK* featured a mean PNmax value (the measure of rhythmicity) of 22.12, while the mean PNmax value of its most abundant genes was 9.22. Only three functions within the nitrogen metabolism (*gdhA*, *nirB,* and *nifD*), one within the carbon metabolism (*ccoN*), and one within the iron metabolism (*isiA*) were less synchronous and rhythmic than the means of their most abundant genes.

**Figure 4.**
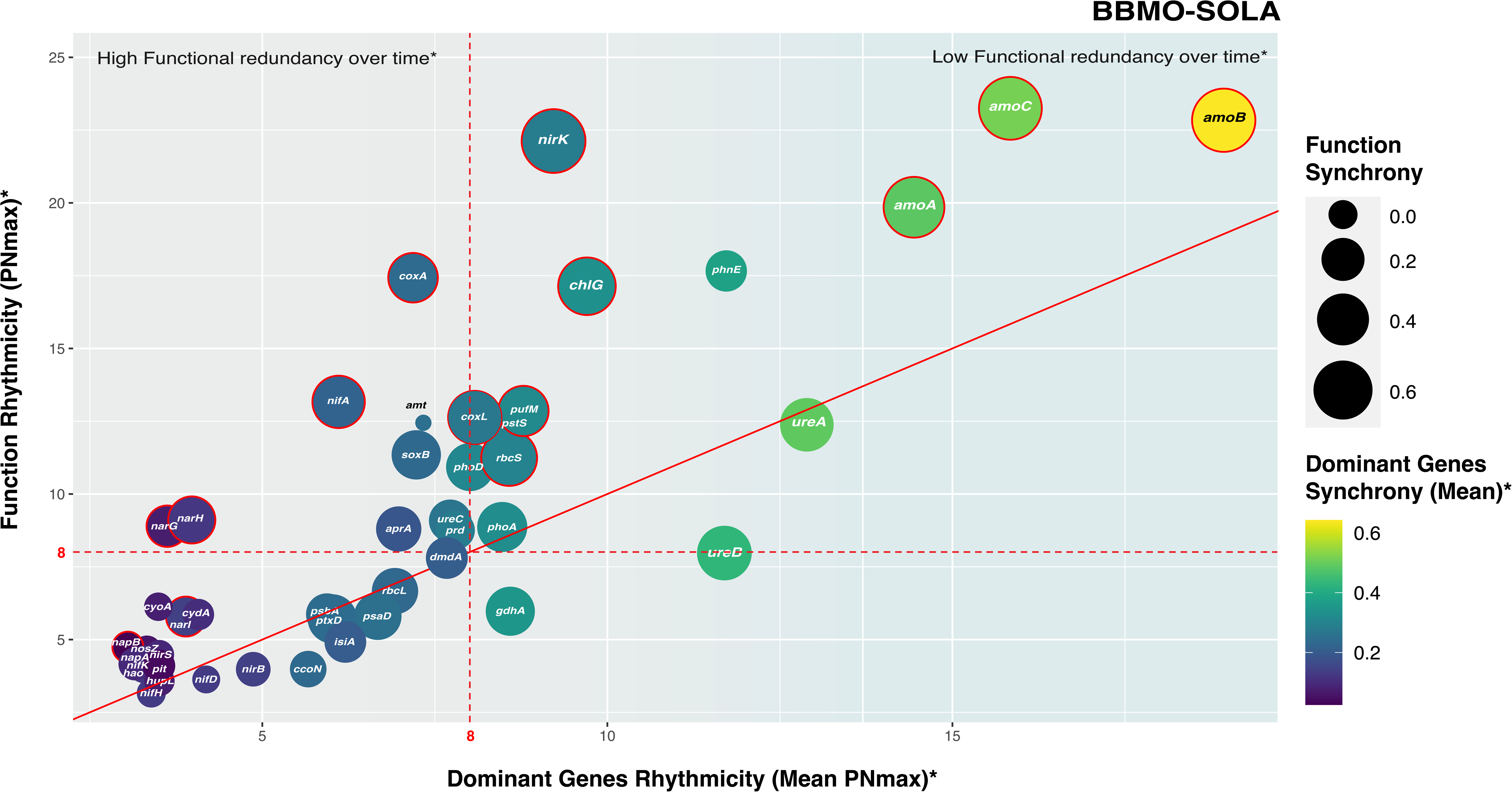
Rhythmicity (y-axis) of biogeochemically key functions in BBMO and SOLA as a function of the rhythmicity of their dominant genes (x-axis). The lines at PNmax = 8 indicate the threshold from which a function or gene is considered rhythmic. The bubble size indicates the synchrony of the biogeochemically key functions, while the synchrony of its dominant genes is shown with a colour gradient. (*) Dominant genes are those that account for more than 70% of the total abundance of a function. Red circles indicate functions showing both higher rhythmicity and synchrony than that of the dominant genes within the function. Note that the synchrony of functions and dominant genes tends to increase with the increasing rhythmicity of functions and genes. Also, observe that functions on the left side of the X-axis (low dominant gene rhythmicity) are candidates for higher functional redundancy and vice versa.

Overall, the preceding results lead to two hypothetical scenarios that may coexist across different functions. In the first scenario, of “high functional redundancy”, compensatory changes in the abundance of functionally redundant taxa lead to the maintenance of the synchrony and rhythmicity of functions at the community level (**Figure 5A, left**). This is exemplified by the *narH* function, which encodes a subunit of the nitrate reductase enzyme (Fig. 5B, left). As depicted, while the community changes monthly with an elevated turnover rate (0.82), the function dynamics prevail with a high rhythmicity. In turn, in a scenario of “low functional redundancy”, the synchrony and rhythmicity of functions are primarily driven by abundant species (**Figure 5A, right**). In this case, a decrease in the abundance of one species contributing the majority of ORFs to one function will affect the synchrony and rhythmicity of such function, as no other species are expected to increase in abundance to compensate for the reduction in ORFs (**Figure 5B, right**). This scenario is depicted by the *amoB* function, which encodes the methane/ammonia monooxygenase subunit B (Fig. 5B). Here, in concordance with the lower turnover rate (0.55), the function is dominated by a few taxa, especially the genus *Nitrosopumilus,* setting the rhythmic dynamic of the function.

**Figure 5.**
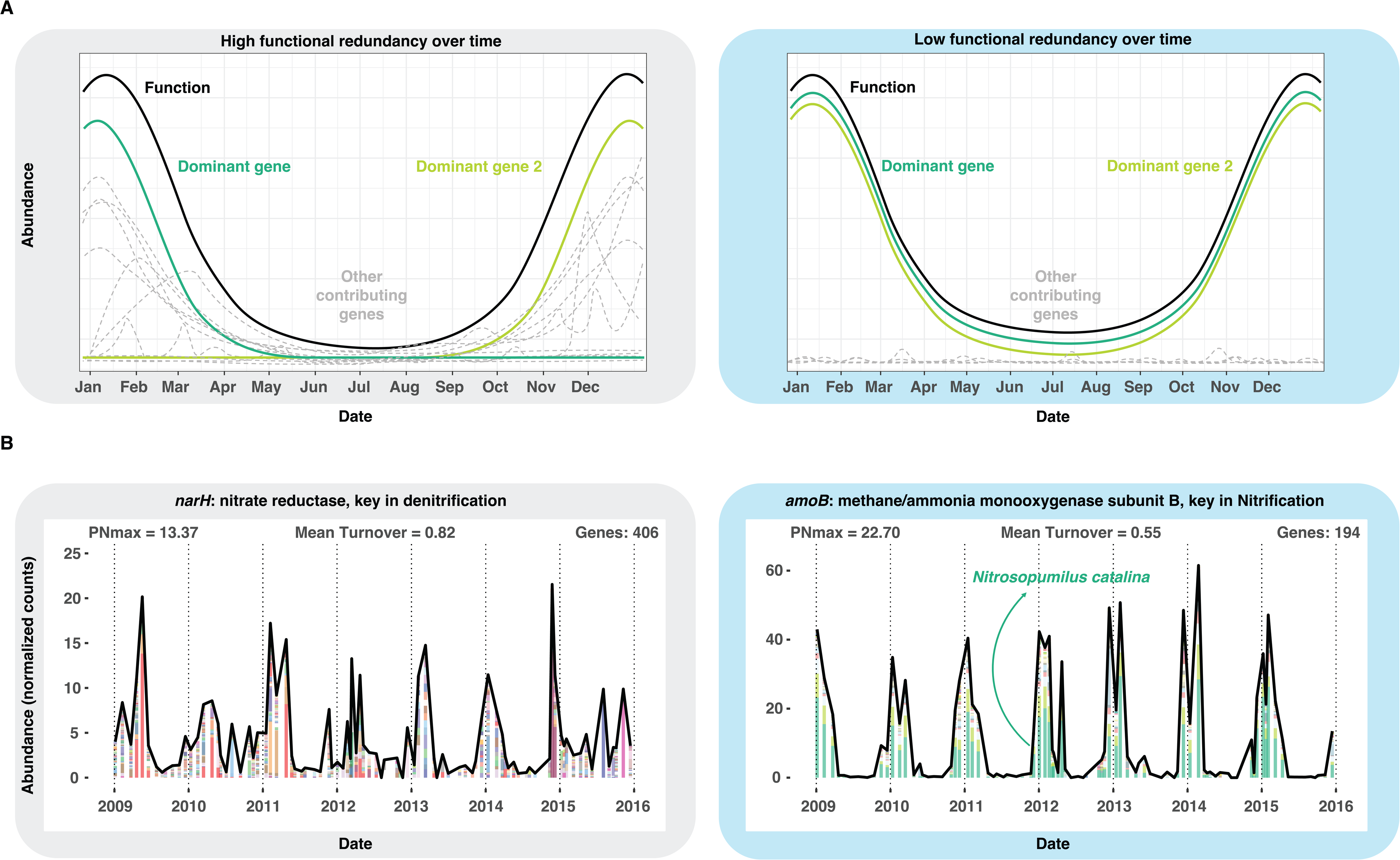
Relationship between functional redundancy and the community dynamics of biogeochemical functions. Panel. **A.** Conceptual diagram displaying the annual abundance of a function (in black), its most abundant contributing genes (in green, termed "dominant genes"), and the remaining less abundant contributors (in grey). In the first scenario (**left**), many genes contribute substantially across the year and dominance shifts frequently, indicating high functional redundancy. In the second scenario (**right**), only a few genes consistently dominate over time, with minimal contribution from others, illustrating low functional redundancy, where the function depends on a small, stable set of genes. **Panel B, left.** Temporal abundance distribution of the *narH* function showing a rhythmic pattern (PnMax = 13.37) in SOLA; colored bars represent the contribution of different genes (ORFs) to the overall function abundance. Note that from the total of 406 genes annotated to this function, as indicated in the figure, there is a monthly change of the contributing genes as depicted by the mean turnover. **Panel B, right.** Temporal distribution of the *amoB* function showing a rhythmic pattern (PnMax = 22.70) in SOLA; colored bars represent the contribution of different genes (ORFs) to the overall function abundance. Note that from the 194 genes annotated to this function, only a few dominate in abundance. As a consequence, the contributing community changes little each month, as indicated by the mean turnover at the top.

Ultimately, remarkable differences in the functional redundancy patterns were observed between BBMO and SOLA microbiomes for the selected 45 biogeochemical functions (**Figure S4)**. Functions in SOLA displayed a higher (function rhythmicity)/(dominant genes rhythmicity) ratio compared to their BBMO homologs (note the upward and leftward scrolling of almost all functions in **Figure S4**). This suggests that, for several functions, SOLA exhibits higher rhythmicity than BBMO, even when the most abundant ORFs displayed non-rhythmic dynamics, emphasizing the ‘high functional redundancy’ scenario for SOLA.

## DISCUSSION

Gaining insight into the mechanisms that shape the structure of marine microbiomes and their potential responses to ongoing global change is crucial. Functional redundancy is widely regarded as a critical feature that supports the stability and resilience of ocean microbiomes. Although its prevalence and significance are still under debate (*24*), investigating how it manifests in environmentally similar and interconnected microbial communities may offer critical insights into its role in shaping community structure and driving emergent ecological dynamics. Patterns of community synchrony and rhythmicity can further elucidate the extent to which functional redundancy contributes to the structure of microbial communities. However, the extent to which marine microbiomes interconnected by oceanic currents exhibit synchronous or unsynchronous dynamics over extended periods, as well as the influence of functional redundancy in shaping these patterns, remains unclear (*29*, *46*). Here, we conducted a long-term analysis of two neighboring coastal marine microbiomes in the Mediterranean Sea, which are linked via ocean currents and subject to rather comparable environmental fluctuations. Our objective was to ascertain the degree of rhythmicity and synchrony in their monthly ecological dynamics across 7 years (2009–2015) and to determine the links between the observed dynamics and functional redundancy. We found that the microbiomes in BBMO and SOLA are highly rhythmic at the three organizational levels we investigated: metabolic functions, genes (ORFs), and OTUs. This aligns with previous findings on the seasonality of these temperate marine microbial ecosystems (*12*, *44*, *47*). Environmental variables in both locations, such as temperature, sunlight, and nutrient availability, undergo seasonal variations (*20*, *48*), and microorganisms often track these changes, leading to rhythmic seasonal dynamics. However, despite the significant correlation between rhythmicity and synchrony (Pearson correlation coefficient = 0.51, p-value <0.001, R^2^= 0.26), our analysis unveiled low synchrony between the microbiomes at both locations across the three organizational levels investigated. This suggests that although both communities respond rhythmically to seasonality, the timing of their responses is not strictly synchronous, reflecting the complexity of microbial community dynamics in the ocean. Thus, our results point towards microbial dynamics that are rhythmic but, at the same time, highly idiosyncratic. This aligns with observations in other microbiomes, such as the reported remarkable singularity of mammalian-associated microbiomes (*34*, *49*), referring to their unique compositions shaped by individual-specific factors, including human microbiomes (*50*). Despite its relevance, this phenomenon has received limited attention in marine environments, likely due to the traditional focus on broad-scale commonalities rather than site-specific long-term dynamics. For example, recent global studies (*51*, *52*), have explored the ecological mechanisms that structure the ocean microbiome, whereas site-specific temporal dynamics have received comparatively less attention. On the other hand, studies like Ottesen *et al.* (*29*) have investigated short-term synchrony in gene expression among sympatric marine microbial populations over diel cycles. However, these approaches do not capture longer-term, site-specific dynamics. As more marine long-term studies emerge (e.g., BBMO, SOLA), such individualistic patterns are beginning to surface. This highlights the complexity of microbiomes in the ocean, where, despite physical connectivity facilitated by currents (*18*), intrinsic factors might dominate over dispersal, leading to specific community compositions and functions.

High idiosyncrasy in microbial dynamics may suggest that these communities are highly adaptable to the local environmental conditions, which may contribute to their resilience and stability in changing environments (*53*). Additionally, it may suggest that the strength of environmental selection is at least moderate, sufficient to drive seasonal patterns in both microbiomes, yet not strong enough to impose a uniform response in both communities. Strong selective pressure (such as in extreme habitats) is expected to lead to the dominance of certain species or traits well-adapted to specific environmental conditions, displaying recurrent patterns over time and across communities. If two connected microbiomes experience strong and similar selective pressures, one expectation is to observe a high degree of synchrony in their dynamics (*7*, *8*). The similar environmental heterogeneity, coupled with the low synchrony observed in BBMO and SOLA microbiomes, suggests moderate environmental selection. In sum, despite the apparent similarities in environmental variation between the two neighboring localities and their presumably high connectivity, the selective pressures promoting synchrony between the microbiomes were likely not strong enough to overcome the idiosyncrasy of each community. Thus, BBMO and SOLA microbiomes retain distinctive characteristics, suggesting that other ocean microbiomes may exhibit similar site-specific dynamics and prompting questions about the boundaries of marine microbial communities.

A moderate strength of environmental selection enables a variety of species to coexist, many of which can perform overlapping functions, a phenomenon known as functional redundancy (*23*). The latter is expected to decrease the species synchrony between communities since multiple ecologically equivalent taxa respond differently to environmental variations while carrying out the functions needed by the community. Although several microbial taxonomic groups are broadly recurrent over time in both time series (*54–56*), the observed limited synchrony points to significant functional redundancy within both microbiomes. Our findings align with previous studies suggesting significant functional redundancy in the global ocean microbiome, particularly within broad functional categories (*24*). This is coherent with the results from sunlit global-ocean gene catalogues (*32*). In the previous study, when comparing taxonomic and functional variability in the ocean, a high taxonomic diversity was found, even at the phylum level, while gene abundances, when grouped into functional categories, remained relatively stable. Similarly, Louca *et al.* (*57*) compared taxonomic and predicted functional profiles across the global ocean and observed high taxonomic richness and variability within functional groups, further supporting this pattern. In turn, an earlier study from SOLA showed that taxonomy and gene content were highly correlated over time (*35*), suggesting little redundancy at the community level. These contrasting results emphasize the importance of the *redundancy* definition: strict redundancy, which includes the complete set of functions of an organism (species can fully replace each other), versus partial redundancy, involving only specific functions or traits. The functional level chosen for redundancy studies is also important (gene [ORF], functional category, or community) (*35*).

Interestingly, despite the low synchrony observed in BBMO and SOLA, no apparent differences in synchrony were found between broader (metabolic functions) and finer levels, such as genes (ORFs) and OTUs. As previously hypothesized, high functional redundancy could be depicted by cases where metabolic functions exhibit synchronous dynamics over time, while the genes (ORFs) or taxa contributing to those do not (*44*). Ecologically-relevant metabolic functions are often strongly coupled to specific environmental factors (*2*, *58–60*). However, they do not necessarily appear coupled in time to particular taxonomic assemblages (*21*, *61*, *62*). Our results indicate that microbial communities in both locations share specific functions; however, the taxa performing these functions may occupy slightly different ecological niches, shaped by site-specific biotic or abiotic environmental factors (partial redundancy). As a consequence, even when performing the same functions, the timing of the taxa distributions may vary between microbiomes, resulting in site-specific temporal offsets that lower function-level synchrony.

This pattern becomes more evident when examined in a site-specific context. In the SOLA microbiome, our analysis revealed higher rhythmicity at the level of function than at the lower organization levels of gene (ORF) and OTU, pointing to a high functional redundancy scenario. When multiple microbial taxa contribute to the same metabolic processes, the arrhythmic behaviors of their genes (ORFs) and OTUs may combine, resulting in a certain degree of rhythmicity in the overall function (*44*). This pattern of rhythmicity was not observed in BBMO, which highlights the fine-grained differences between both sites and could also be an explanation for the displayed low synchrony at the three levels investigated. Although the heterogeneity of the analyzed environmental variables was similar at both locations, the coefficient of variation of each measured environmental parameter was significantly higher in SOLA than in BBMO. The coefficient of variation reflects the extent of variability in the recorded environmental values. Here, they likely represent the impact of occasional riverine flash flood events in the vicinity of SOLA, contributing to increased environmental variability, particularly through sporadic reductions in salinity (*20*). Additionally, local wind regimes, such as the Tramontana, which is known to affect the SOLA area but not BBMO, may contribute to this variability by resuspending sediments and influencing nutrient dynamics (*63*). Microorganisms are highly sensitive to sudden environmental perturbations, and even small changes in variables such as salinity or nutrient availability (*21*) can significantly affect community composition. Yet, bulk community function may not necessarily change. It is worth noting that because individual functions can exhibit distinct temporal dynamics, analyzing them collectively may mask specific rhythmic patterns, reinforcing the need for function-level resolution. Building on this, we considered a set of 45 key biogeochemical functions from different metabolic pathways, including carbon, nitrogen, phosphorus, and sulfur metabolisms, H_2_ oxidation, and iron-stress responses. Although they displayed distinct dynamics, a consistent pattern emerged: as a general trend, the 45 key biogeochemical functions exhibited higher rhythmicity in SOLA than in BBMO. Yet, dominant genes (ORFs) performing SOLA functions were substantially less rhythmic than their BBMO counterparts. That is, the SOLA microbiome was less dominated in terms of ORFs per function than the BBMO microbiome, yet functions were more rhythmic in SOLA than in BBMO. This further supports the notion that SOLA may exhibit higher functional redundancy than BBMO. In SOLA, sudden environmental perturbations may change microbial community structure but not necessarily the ecosystem function.

The idea that diverse and non-necessarily rhythmic microorganisms may contribute to a rhythmic function challenges the ’biomass ratio’ hypothesis. This concept states that the characteristics of an ecosystem depend on both the species’ ecological traits and their respective contributions to the overall biomass of the community (*61*). In essence, it suggests that ecosystem functions are governed primarily by the traits of the most abundant species. Similar perspectives have also been proposed in the context of marine microbiomes (*64*, *65*). However, recent research has shown that rare species can play significant roles in biogeochemical cycles and may act as hidden drivers of microbiome function (*62*). This is the case of methane consumption in riparian wetlands, which is modulated by the rare biosphere through niche partitioning (*66*). In this context, the dynamics and intensity of methane consumption align with the relative abundance and activity of specific rare subgroups of methane-oxidizing bacteria. Other studies employing stable isotopes to assign species-specific roles in major biogeochemical cycles have also emphasized the profound impact of rare microbial species on globally significant processes (*67–69*).

Here, we observed a comparable phenomenon for the *narH* function. The *narH* encodes one of the subunits of the nitrate reductase enzyme, which is involved in the conversion of nitrate (NO₃⁻) to nitrite (NO₂⁻) during the process of denitrification (*70*). Denitrification is a crucial microbial metabolic pathway in marine ecosystems, especially in environments where oxygen availability is limited. This process is important in the nitrogen cycle, as it helps regulate nitrogen availability in marine ecosystems and affects nutrient cycling. This is typically a highly biodiverse function that can be performed both by bacteria, including SAR11, and by archaea (*71*). Over the 7-year analysis period, we observed substantial monthly variation in the taxa contributing to *narH* in SOLA. Rare, or subdominant contributors, that is, taxa accounting for less than 70% of the total function abundance, were identified, exhibiting non-rhythmic dynamics. As a result, the rate at which species were replaced over time was remarkably high, with a mean turnover rate of 0.82. However, the overall dynamics of the *narH* function remained rhythmic (PNmax = 13.37), with most of the abundance occurring during spring-winter. The leaky nature of denitrification intermediates, combined with high taxonomic turnover and functional redundancy, suggested a distributed maintenance strategy, where multiple taxa may transiently contribute to a costly function whose benefits are diffusely shared across the microbial community. The opposite scenario was also observed. This is the case of *amoB*, one of the enzyme subunits involved in obtaining nitrogen from ammonia (NH_4_^+^) during the nitrification process (*70*) and a key step in transforming nitrogen compounds in marine environments. Here, the taxonomic community contributing to *amoB* in SOLA remained constant throughout the 7 years, with a clear dominance of the abundant archaeal class Nitrososphaera, specifically the genus *Nitrosopumilus*. In this case, the rhythmicity of the function was mostly driven by the dominant taxa. This is consistent with the literature since *amoB* is a less biodiverse function known to be mainly driven by a few archaeal genera (*70*).

Despite the anticipated higher biodiversity of the global ocean (*Tara*), the taxonomic composition at the class level for the 45 key biogeochemical functions in BBMO and SOLA followed a similar pattern to that observed at a global scale. For example, the aforementioned *narH* function was carried out in BBMO and SOLA by diverse taxonomic groups, including Actinobacteria, Alphaproteobacteria, Gammaproteobacteria, and Nitrospinia. In the global ocean, the same groups were identified, except for the absence of Actinobacteria and the inclusion of Marinisomatia. For both BBMO and SOLA, as well as the global ocean, *narH* displayed high species evenness (low dominance) (*72*). Furthermore, for the *amoB*, identical taxonomic groups were identified in BBMO, SOLA, and the global ocean (Nitrosophaeria and Gammaproteobacteria). Additionally, low species evenness was observed in all cases, emphasizing the dominance of a few species in performing this function at a global scale. In other functions, including *psbA*, *phoA*, *phoD*, *nirB*, *cydA*, *soxB*, *coxA*, *amt*, and *prd,* the elevated Pearson correlation between the relative abundance profiles of taxonomic classes per function across BBMO, SOLA, and *Tara* also indicated the same pattern of similarity between both time series and the global ocean. In summary, the similar taxonomic composition and species evenness observed between BBMO, SOLA, and the global ocean for the 45 key biogeochemical functions analyzed suggest that the patterns identified in the two studied time series reflect broader, widespread trends in marine ecosystems.

Our analysis revealed that both rhythmic and non-rhythmic taxa contributed to the recurrent functional dynamics observed in both microbiomes. This can be partially explained by ‘functional synergy,’ which, in this context, refers to the dynamics or patterns observed at a higher organizational level (functions) that arise from the interplay of multiple dynamics occurring at a lower organizational level (taxa or genes) (*23*). The closest example of this observed phenomenon in our study is the dynamics of the B_12_-related metabolism in SOLA. Here, non-rhythmic genes, such as *cobF*, *cobG*, and *cobA-btuR*, contributed to the rhythmic dynamics of B_12_ biosynthesis (*44*). Functional synergy may encompass self-organization processes among microbial species, where their combined metabolic activity emerges from a complex interplay of dynamics (*73*). Similarly, as observed in polymicrobial infections, the combined impact of multiple microorganisms on disease outcomes can be more severe than the effects of individual microbes alone (*74*). In marine microbiomes, similar interactions may enhance metabolic efficiency through functional redundancy, where multiple species perform overlapping functions, but also contribute to metabolic complementarity, where different taxa support each other’s metabolism; for example, one taxon producing a by-product that another will use (*23*, *44*). While our study does not directly test these mechanisms, the patterns we observed may, to some extent, reflect underlying microbial interactions that altogether sustain functional stability despite taxonomic fluctuations. Investigating whether these dynamics arise from cooperative metabolic exchanges or niche differentiation remains an important avenue for future research.

## CONCLUSION

The comparative study of two connected long-term marine microbial observatories revealed a significant idiosyncrasy in marine microbiome dynamics depicted by the low synchrony observed across different organizational levels, including functions, genes, and taxa, despite comparable environmental heterogeneity. However, both microbiomes exhibited rhythmicity across all organizational levels, with particularly high rhythmicity observed at the functional level, especially in SOLA, where functions were more rhythmic than the genes or taxa contributing to them. This particularity suggests substantial functional redundancy in both microbiomes, a characteristic that may also apply to other microbiomes over extended time periods. The analysis of 45 key biogeochemical functions revealed high synchrony and rhythmicity in certain functions, even when the most abundant contributing genes exhibited low synchrony and rhythmicity, suggesting both functional redundancy and synergy in critical metabolisms, potentially reflecting self-organization processes. Our findings suggest that the observed trends may be common across marine environments, given the similar patterns identified in the global ocean. Lastly, the results advance our understanding of how the ocean microbiome may respond to global change by shedding light on the roles of functional redundancy and self-organization in shaping community functions over time under varying environmental conditions.

## MATERIALS AND METHODS

### Sampling, DNA extraction, and shotgun sequencing

Coastal surface water samples (0.5 m depth) were collected at the Blanes Bay Microbial Observatory (BBMO, 41°40′N, 2°48′E; http://bbmo.icm.csic.es) in the Bay of Blanes, Catalonia, Spain. BBMO is an oligotrophic coastal site located about 1 km offshore, with a depth of approximately 20 meters and minimal influence from rivers or human activity. Furthermore, additional samples were collected at the Banyuls Bay Microbial Observatory (SOLA, 42°29’18.2"N 3°08’35.6"E) in the Bay of Banyuls-sur-Mer, Northern Catalonia region, France. SOLA is also an oligotrophic coastal site, situated about 500 m off the coast of Banyuls-sur-Mer, with a depth of around 26 meters and limited human impact (*75*). However, unlike BBMO, SOLA experiences sporadic winter storms, which resuspend nutrients from the sediments into the water column. Additionally, flash floods from nearby rivers further enrich SOLA’s waters with nutrients (*76*). The stations are separated by ∼150 km in the Northwestern Mediterranean Sea and are connected by a dominant south-western marine current, also known as the Liguro-Provençal-Catalan Current. This current travels along the Italian coast west of Genoa, as well as the French and Catalan coastlines, with surface speeds that can reach 1 m/s (*77*). Despite being a stable feature, in the summer months the current loses strength to the detriment of coastal water masses flowing from south to north of the Iberian peninsula, potentially increasing the biotic and abiotic differentiation between BBMO and SOLA (*18*).

In BBMO, 6 L of 200-μm pre-filtered surface seawater were sequentially filtered through a 20-μm mesh, a 3-μm pore-size polycarbonate filter (Poretics), and a 0.22-μm Sterivex Millipore (Merck-Millipore) filter using a peristaltic pump. At SOLA, a subsample of 5 L from 10 L Niskin bottles was sequentially filtered through 3 μm pore-size polycarbonate filters (Merck-Millipore, Darmstadt, Germany) and 0.22-μm Sterivex Millipore (Merck-Millipore). Sterivex cartridges containing the pico-fraction (0.22-3 μm) of the microbial biomass were stored at –80 °C until nucleic acid extraction for both sampling stations. Note that sampling at both locations was part of the regular sampling at both time series. Monthly samples from January 2009 to December 2015 (7 years) were compiled, generating a dataset of 84 samples for the BBMO station and 90 samples for the SOLA station. While SOLA featured more than one sample in certain months, the BBMO dataset was limited to one sample per month.

DNA extractions of the BBMO samples were performed following the protocol described by Schauer et al., (*78*) in which a lysozyme solution (20 mg/mL) was added to the Sterivex cartridges to lyse cells. Subsequently, a second incubation with proteinase K (20 mg/mL) was applied, and the pooled lysates were then extracted twice with an equal amount of phenol-chloroform-isoamyl alcohol (25:24:1, pH 8) and once with an equal amount of chloroform-isoamyl alcohol (24:1). Finally, a purification in Amicon units (Millipore) was performed. DNA extractions for the SOLA samples were performed following the protocol described by Hugoni et al. (*79*) using a lysozyme solution (20 mg/mL) for cell lysis with a second incubation with proteinase K (20 mg/mL), and the pooled lysates were then extracted using the AllPrep DNA/RNA kit (Qiagen, Hilden, Germany). Sampling and extraction protocols do not generate discrepancies and remain comparable across studies (*80*).

Before sequencing the BBMO metagenomes, DNA quality control was performed using an agarose gel, a Qubit fluorometer, and a Nanodrop. From January 2009 to December 2011 (3 years), the samples were sequenced on an Illumina HiSeq4000 (2 x 150 bp) platform, and from January 2012 to December 2015 (4 years) on an Illumina NovaSeq6000 (2 x 150 bp) at the Centre Nacional d’Anàlisi Genòmica CNAG, Barcelona, Spain. About 22.5 billion reads were produced. For sequencing SOLA metagenomes, DNA quality control was performed using the Agilent High Sensitivity kit (Agilent Technologies, Santa Clara, CA, USA). From January 2009 to December 2011 (3 years), the samples were sequenced on an Illumina NovaSeq6000 platform (2 x 150 bp) at the Centre Nacional d’Anàlisi Genòmica (CNAG), Barcelona, Spain. From January 2012 to February 2015 (∼3 years), the samples were sequenced on an Illumina HiSeq2500 platform (2 × 100 bp) at GenoScreen, Lille, France, and from March 2015 to December 2015, the samples were sequenced on an Illumina NovaSeq6000 (2 x 150 bp) at CNAG. About 19 billion reads were produced for SOLA. In total, ∼ 41 billion reads were produced for BBMO and SOLA.

### Bioinformatics analyses

All reads were trimmed with CUTADAPT v1.16 (*81*) to remove adapters and low-quality reads. Each sample was then assembled individually with MEGAHIT v1.1.3 (*82*) using the meta-large presets. Contigs were checked using QUAST (*83*). Genes (ORFs) were predicted in the contigs from each sample using Prodigal v2.6.3 (*84*) and MetaGeneMark v3.38 (*85*). The predicted genes from the different samples were pooled and dereplicated by clustering them at 95% identity and 90% coverage of the shorter ORF using Linclust v10 (*86*). After that, a gene catalogue of 375,380,895 genes was generated for the BBMO-SOLA dataset, of which 209,195,684 genes were longer than 250 bp (*87*).

Predicted genes were annotated using Diamond blastp 0.9.22 (*88*) for the Carbohydrate-Active enzymes database (CAZy) (*89*) and the Kyoto Encyclopedia of Genes and Genomes (KEGG) (*90*), Rpsblast 2.7.1 (*91*) for the Clusters of Orthologous Genes database (COG) (*92*), HMMER3 for the evolutionary gene genealogy Non-supervised Orthologous Groups (eggNOG) (*93*), and the Protein families database (Pfam) (*94*). In total, 18,625,274 genes were annotated with CAZy (13.87%), 48,838,588 genes with KEGG (36.38%), 74,954,838 genes with COG (55.83%), 73,345,901 genes with Pfam (54.63%), and 86,545,308 genes with eggNOG (64.46%).

Gene abundances per sample were calculated by mapping metagenome reads back to the catalogue using BWA v0.7.17 (*95*), and obtaining the number of counts per gene using HTSeq v0.10.0 (*96*). Gene counts were normalized within and among samples by gene length and metagenome size. Normalized gene abundance tables were generated, including the abundance of each gene (ORF) in each sample. From these, we calculated the corresponding functional abundance tables by adding all the normalized abundances of all genes that were annotated to a specific function within a given database (i.e., CAZy, KEGG, COG, eggNOG, Pfam) (**Figure S5**).

The taxonomic community composition of the different samples (metagenomes) was determined using mTAGs (*97*). This approach uses a reference database based on the clustering of sequences from the full-length SILVA SSU database version 138 (*98*) into Operational Taxonomic Units (OTUs) at 97% identity. Metagenomic reads that are identified to belong to the rRNA are then mapped against the reference OTUs, and their abundances are estimated. Functional genes were taxonomically annotated against the Genome Taxonomy Database (GTDB) release 95 (*99*) using MMSeqs2 v11-e1a1c with a sensitivity value of 5.7 (*100*) and specifying ranks at which the Lowest Common Ancestor (LCA) should be determined. Taxonomic annotations were reported primarily at the class level for consistency, given that resolution from metagenomic data was limited by database coverage and gene conservation. In a few cases, deeper taxonomic assignments were included when functional markers were highly specific and taxonomic identity was unambiguous.

### Statistics

Data analyses were performed using Python v3.8.5 (*101*) and R v3.5 (*102*). Specifically, functions implemented in the packages *tidyverse* v1.3 (*103*), *vegan* (*104*), *pvclust* (*105*) were used to process the data. All data visualization was performed with *ggplot2* v3.2. In order to calculate the temporal synchrony between function, gene, and OTU abundances in both microbiomes, the *codyn* package was used. The synchrony metric compares the average correlation of each feature with the rest of the aggregated community (*26*). The synchrony function returns a numeric value oscillating between -1 (asynchrony) and 1 (full synchrony). Synchrony values were classified as high (Sy>=0.7), moderate (0.4=<Sy<0.7), and low (0=<Sy<0.4) based on Gross et al (*26*). The rhythmicity of functions, genes, and OTUs was computed using the *lomb* package, which generates the Lomb Scargle periodogram (LSP). LSP is an algorithm based on the Fourier mathematical transformation (*106*, *107*) previously implemented in the study of time series in ecology (*20*, *108*). The peak normalized power (PNmax) of each LSP was calculated and considered rhythmic when PNmax ≥ 8. This threshold was based on previous studies (*20*, *44*).

Data analyses were divided into data- and hypothesis-driven approaches. To avoid biases, given that many genes were only sporadically present and in small quantities, we considered only genes and OTUs with at least 30% occurrence through the dataset. For the data-driven approach, we calculated the synchrony of 429 functions from CAZy, 8,455 functions from KEGG, 4,872 functions from COG, 93,303 functions from eggNOG, and 16,624 functions from Pfam within BBMO-SOLA. In addition, we calculated the synchrony of the selected 6,375,074 genes (ORFs) and 11,499 OTUs. In parallel, the rhythmicity was calculated for 428 CAZy, 8,414 KEGG, 4,872 COG, 96,603 eggNOG, and 16,605 functions from Pfam for BBMO and for 428 CAZy, 8,382 KEGG, 4,872 COG, 96,603 eggNOG, and 16,595 functions from Pfam for SOLA. In addition, rhythmicity was calculated for the selected 5,047,243 genes and 11,402 OTUs from BBMO, and 5,047,240 genes and 11,402 OTUs from SOLA. Environmental heterogeneity was calculated by computing the average dissimilarity between sites based on eight abiotic and biotic variables: Temperature, salinity, day length (h), chlorophyll, NH_4_, NO_3_^-^, NO_2_^-^, and PO_4_^3-^ concentrations. We computed an Euclidean distance matrix for each site and calculated the dissimilarity between sites (Ed) as described before (*109*, *110*). In addition, we calculated the coefficient of variation (CV) for each variable as the standard deviation divided by the mean of each variable. Each variable was converted to z-scores to standardize the data.

For the hypothesis-driven approach, a specific subset of 45 key biogeochemical functions was analyzed as they play an important role in the ocean’s main biogeochemical cycles and can be considered marker genes for these functions. Specifically, for the carbon metabolism, we included *coxL, cyoA, cydA*, *ccoN*, *coxA*, *rbcL, rbcS, pufM*, *chlG*, *prd*, *psaD*, and *psbA*. For the nitrogen metabolism, we included *nosZ*, *nifH*, *nifD*, *nifA*, *nifK*, *napA*, *napB*, *nirS*, *nirB*, *narH, narG*, *narI*, *gdhA*, *amoA*, *amoB*, *amoC*, *nirK*, *ureA*, *ureB*, *ureC, amt*, and *hao.* For the sulfur metabolism, we considered *dmdA*, *soxB*, and *aprA*. For the phosphorus metabolism, we included *pstS*, *pit*, *phnE*, *phoA*, *phoD*, and *ptxD*. Finally, we also considered *isiA* as iron metabolism related and *hupL* for its role in H_2_ oxidation. Further details on the selected functions can be found in the Supplementary Materials.

Cleaning was required for the *coxL* function. In this study, only the aerobic type I Carbon monoxide dehydrogenase (CODH) *coxL* was considered since it is the key enzyme in aerobic CO oxidation by carboxydotrophic bacteria (*111–113*). Given the proximity of CODH with other molybdenum-containing enzymes within the same protein family (*111*), distributed in all domains of life (*114*), positive hits were further analyzed to remove false positives by phylogenetic placements. Putative CODH peptides were removed if they did not cluster with a form I CODH large subunit in a reference database, as explained in the Supplementary Materials.

For each of the selected 45 functions, we calculated Pielou’s evenness index to evaluate how evenly the contributing taxa shared the function’s total abundance (*115*). Values closer to 1 indicate that taxa are perfectly evenly distributed within the function. Namely, all taxa contribute equally. Values closer to 0 mean that the taxa are highly unevenly distributed, meaning one or a few taxa dominate while others are rare. Functions were hierarchically clustered based on Euclidean distances at the Class level of taxonomy. The clustering’s statistical significance was indicated by the p-value, represented as the approximately unbiased (AU) value. The Pearson correlation (PPC) was calculated between BBMO-SOLA and *Tara* Oceans taxonomic distributions of the genes for each function at the class level, using the *stats* R package, as a measure of similarity in taxonomic composition. Among these, dominant genes were defined as those representing >=70% of the abundance of the function in which they were annotated. Turnover between samples was calculated for each of the 45 selected functions as the proportion of taxa (or genes) gained or lost between subsequent samples, relative to the total number of taxa (or genes) observed in both samples. This was performed using the *codyn* R package (*116*).

## ACKNOWLEDGMENTS

We would like to acknowledge all the members of the BBMO sampling team (http://bbmo.icm.csic.es) and the Ecology of Marine Microbes group (http://emm.icm.csic.es) and would like to thank the MARBITS platform of the Institut de Ciencies del Mar (ICM; http://marbits.icm.csic.es) and the Finisterrae III supercomputer at the Centro de Supercomputación de Galicia (CESGA; https://www.cesga.es/) for supporting the bioinformatics analyses. This paper acknowledges the ‘Severo Ochoa Centre of Excellence’ accreditation (CEX2019-000928-S). Views and opinions expressed are however those of the author(s) only and do not necessarily reflect those of the European Union. Neither the European Union nor the granting authority can be held responsible for them.

---

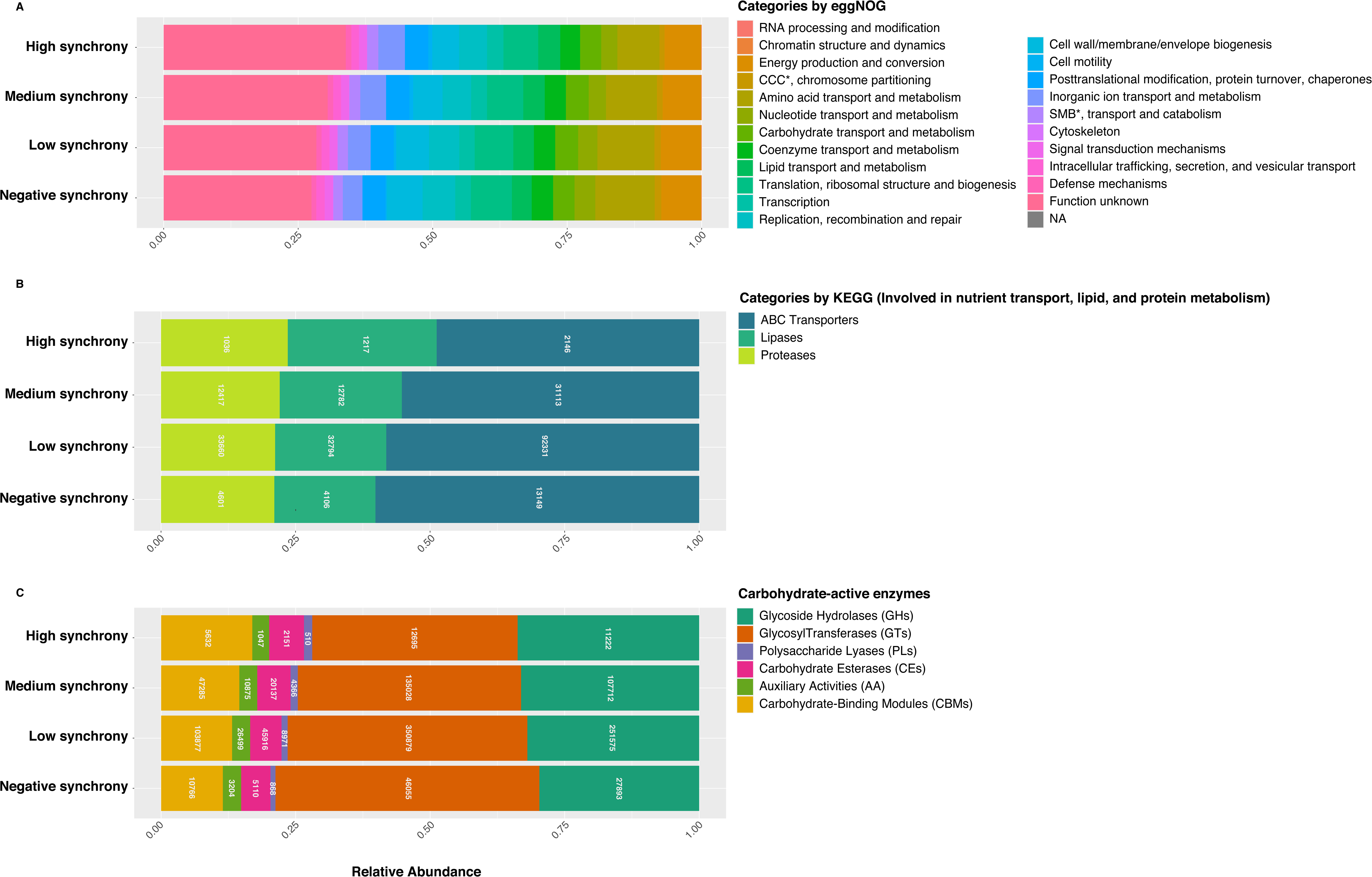

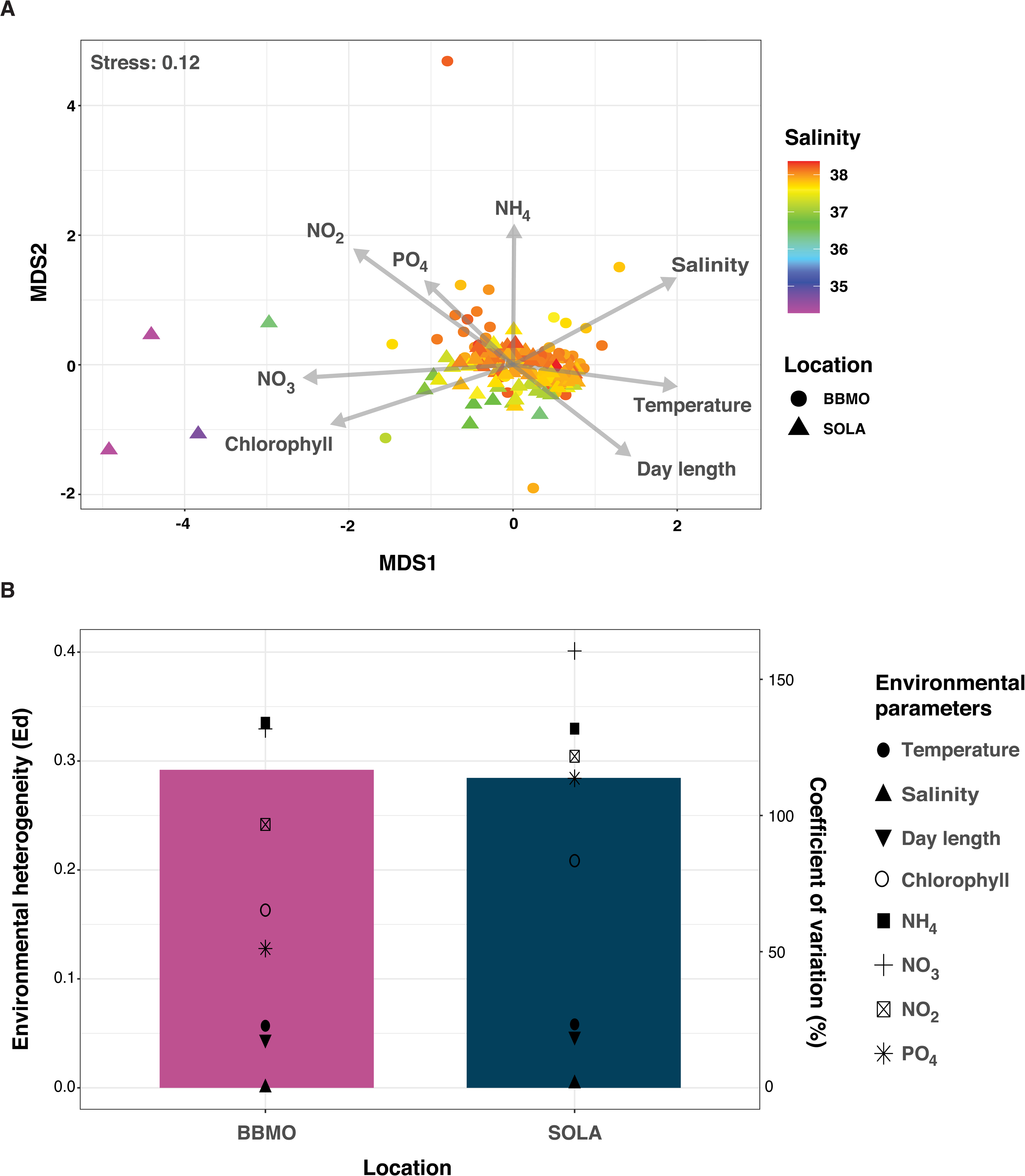

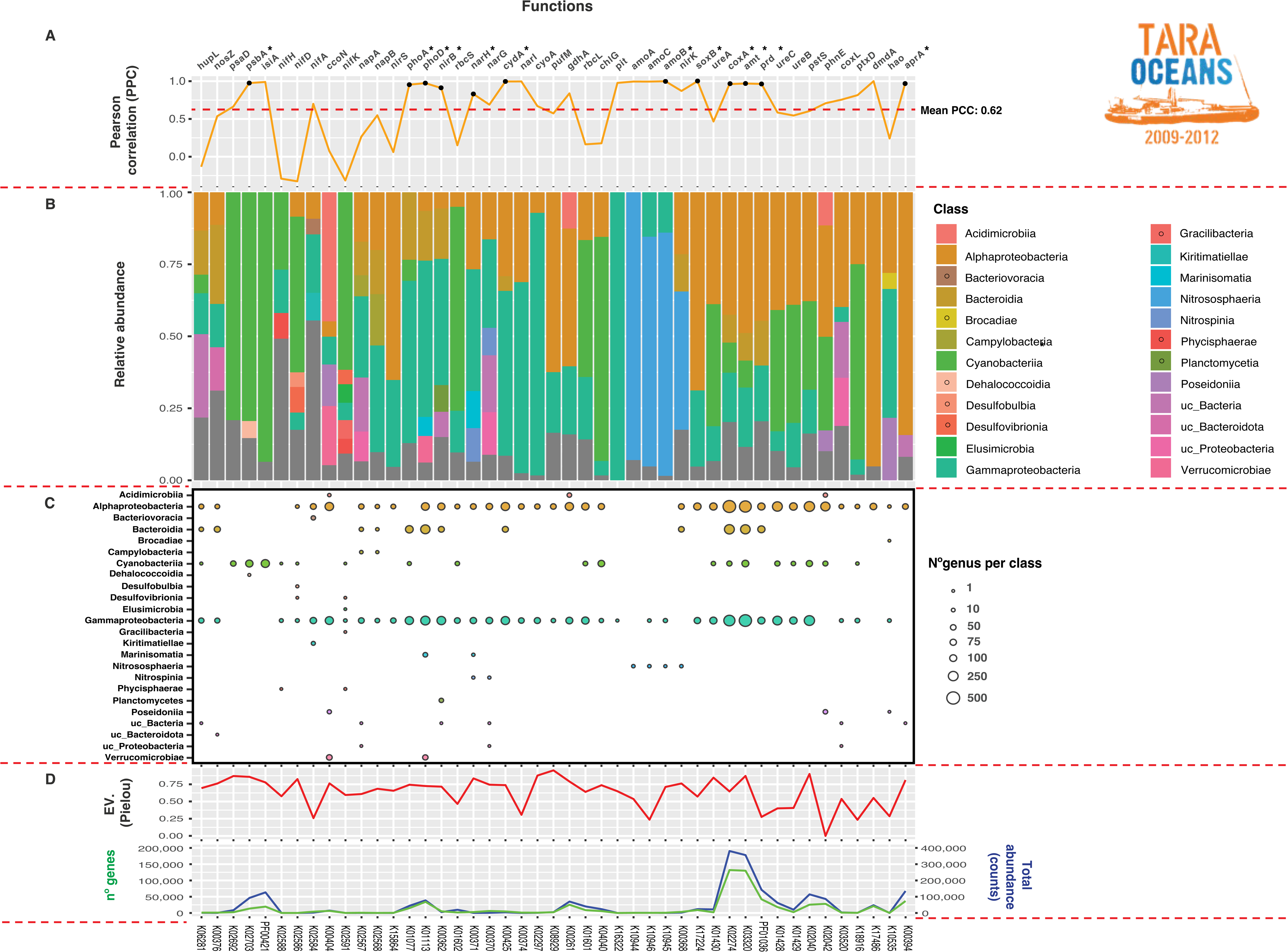

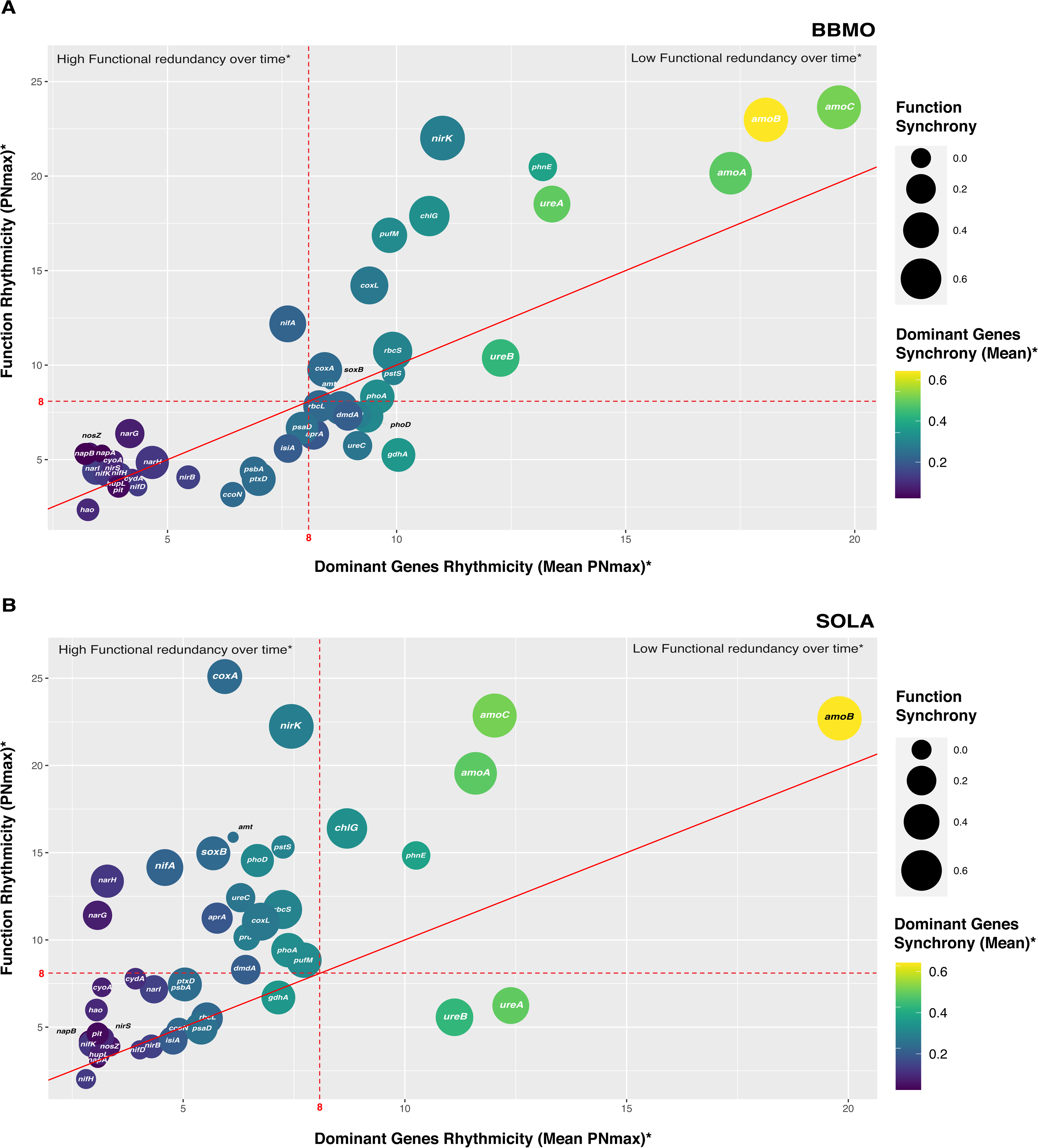

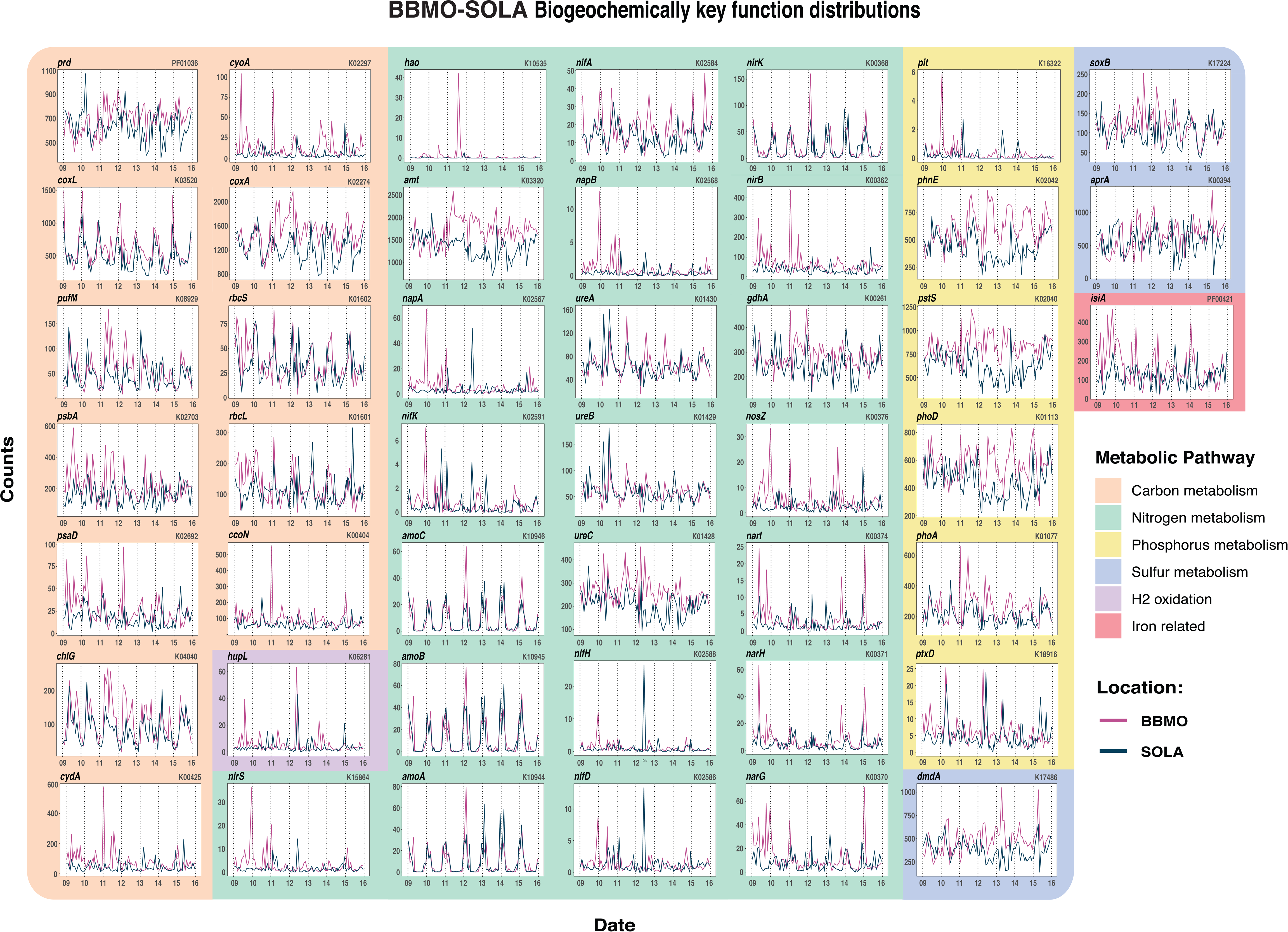

