## Supplementary materials for "Functional redundancy enables emergent metabolic dynamics in marine microbiomes"

### Supplementary Methods

#### Subset of 45 key biogeochemical functions

For the hypothesis-driven approach, a specific subset of 45 key biogeochemical functions was analyzed as they play an important role in the main biogeochemical cycles in the ocean and could be considered marker genes of those functions. The following genes related to carbon metabolism were considered: ***coxL*, *cyoA, cydA, ccoN, coxA, rbcL, pufM, chlG, prd, psaD* and *psbA*.**

*coxL* encodes the carbon monoxide dehydrogenase large subunit enzyme. This gene was selected since heterotrophic bacteria can obtain additional energy from the oxidation of carbon monoxide in the marine environment (*1*). *cyoA* encodes the cytochrome $bo_3$ ubiquinol terminal oxidase in many aerobic bacteria. In *E.coli* it is known to be the component of the aerobic respiratory that predominates when cells are grown at high aeration (*2*). It is reasonable to think that changes in *cyoA* expression are linked to changes in oxygen availability. *cydA*, on the other hand, expresses the cytochrome *bd* ubiquinol oxidase subunit I, involved in aerobic respiration. This enzyme has a high affinity for oxygen and is thought to be important in the survival of bacteria under low oxygen conditions. In contrast to *cyoA*, it predominates when *E.coli* cells are grown at low oxygen concentration (*2*). *ccoN* encodes the cytochrome *c* oxidase *cbb3*-type subunit enzyme I and allows efficient respiration even under low oxygen concentrations (*3*, *4*). *coxA* expresses the cytochrome *c* oxidase *aa3*-type subunit I enzyme, also a component of the respiratory chain that allows for growth in microaerobic conditions (*5*). The genes *rbcL* and *rbcS* are considered marker genes that encode the large and small subunits of the ribulose-bisphosphate carboxylase, included since heterotrophic bacteria are able to fix $CO_2$ through the RuBisCO enzyme (*1*). *pufM* (photosynthetic reaction center M subunit) is a gene marker for the presence of aerobic anoxygenic phototrophic bacteria (AAP) (*1*). *chlG* (chlorophyll/bacteriochlorophyll *a* synthase) is involved in one of the last steps of the biosynthesis of chlorophyll *a*. (*6*). *prd* is included since it encodes the rhodopsin apoprotein as heterotrophic prokaryotic cells can obtain additional energy from sunlight by means of the rhodopsin-based photosystem (*1*). The product of *psaD* (photosystem I subunit II) forms a complex with ferredoxin and ferredoxin-oxidoreductase in photosystem I (PSI) reaction center, also important in the carbon fixation through photosynthesis (*7*). *psbA* expresses the photosystem II P680 reaction center D1 protein, also involved in inorganic carbon fixation (8).

The following functions were analyzed to track the changes in nitrogen metabolism: ***nosZ, nifH, nifD, nifA, nifK, napA, napB, nirS, nirB, narH, nirK, narG, narI, gdhA, amoA, amoB, amoC, ureA, ureB*** **and** ***ureC, amt, and hao***. *nosZ* is a marker gene for denitrification under anaerobic conditions in the presence of nitrate or nitrous oxide (*9*). As some cyanobacteria and heterotrophic bacteria are capable of $N_2$ fixation, *nifH, nifD, and nifK* are the structural nitrogenase genes (*10*), while *nifA* is a regulatory gene (*11*). *napA* and *napB* encode the periplasmic nitrate reductase enzyme essential for dissimilatory nitrate reduction to nitrite (*12*, *13*) and *nirS* (nitrite reductase) for nitrite to nitric oxide (*14*). *nirB* (nitrite reductase large subunit) and *narH* (nitrate reductase, beta subunit) participate in the dissimilatory reduction pathway from nitrite to ammonia (*1*, *15*). *nirK* expresses the nitrite reductase, an enzyme that catalyzes the reduction of nitrite to nitric oxide (*17*). Both *narG* and *narI*, which encode subunits of the nitrate reductase, are also involved in denitrification. *gdhA* (glutamate dehydrogenase) is part of the nitrogen metabolism pathways since it catalyzes the reversible oxidative deamination of glutamate to alpha-ketoglutarate and ammonia (*16*). *amoABC* (methane/ammonia monooxygenase subunit A) are marker genes for ammonia oxidation (*1*). *ureA*, *ureB*, and *ureC* encode the urease subunits gamma, beta, and alpha, respectively, involved in urea transport and degradation to ammonium (*8*), while *amt* is a widespread ammonium transporter gene, a marker for direct ammonium acquisition (*18*). *hao* expresses hydroxylamine dehydrogenase, a periplasmic enzyme that oxidizes hydroxylamine, coming from the oxidation of ammonia by the ammonia monooxygenase (Amo), to nitrite (*19*) and is therefore a relevant marker of nitrification.

As marker genes in sulfur metabolism, the following three were considered: *dmdA*, *soxB* and *aprA*. *dmdA* encodes the dimethylsulfoniopropionate (DMSP) demethylase enzyme, involved in the first reaction in the pathway of organic sulfur assimilation in bacteria via DMSP (*1*). *soxB* (S-sulfosulfanyl-L-cysteine sulfohydrolase) is present in a wide diversity of bacteria and is needed for the oxidation of inorganic S compounds (*16*, *18*) as is *aprA* (dissimilatory adenosine-5′-phosphosulfate reductase) in both aerobic sulfur oxidizers and sulfate reducing bacteria (*20*, *21*).

The phosphorus cycle is represented by the genes *pstS, pit, phnE, ptxD, phoA,* and *phoD*. The product of *pstS* is part of a high-affinity phosphate transport system. Under P-limiting conditions, a phosphate-specific transport system uses ATP-mediated transport that contains a high-affinity phosphate-binding protein (PstS) to meet P needs (*22*). On the other hand, *pit* (low-affinity inorganic phosphate transporter) is typically expressed under high phosphorus concentrations, mainly detected in post-bloom conditions (*18*). *phnE* encodes a phosphonate transport system, permease protein, to obtain phosphorus from dissolved organic phosphorus (*1*). The enzyme PtxD catalyzes the conversion of phosphonates to phosphate, making them available as a phosphorus source for microbial cells (*1*). Lastly both *phoA* and *phoD* (alkaline phosphatase A and D, respectively) are needed to obtain phosphorus from phosphoesters present in the marine dissolved organic phosphorus (*1*).

Iron is a limiting nutrient under certain conditions in the marine environment. Iron deficiency is known to decrease the amount of primary productivity in both marine and freshwater ecosystems. Cyanobacteria remodel the expression of *isiA* (chlorophyll-protein complex CP43) in response to iron deficiency and excess of light (*23*). Finally, the *hupL* (Ni, Fe hydrogenase large subunit) was also included in the analysis since some heterotrophic marine microorganisms use hydrogen gas as an additional energy source (*24*).

***coxL* function filtering**

A reference phylogenetic tree was used to identify form I CODH. Reference genomes were those in Baltar et al. (*25*) and their open reading frames (ORFs) were determined with Prodigal (*26*). To obtain the sequences to construct the reference tree, the corresponding peptides were retrieved from the genome database using TIGR02416 running HMMER3 (*27*) version 3.3.2, with their suggested gathering score. The retrieved peptides were confirmed to have the characteristic signature AYxCSFR at the catalytic site. After removing duplicates, with the resulting peptides, phylogenetic trees were predicted with IQ-TREE (*28*) version 2.1.4-beta.

The metagenome peptides were placed on the reference phylogenetic trees to confirm their functional prediction as form I CODH. To do the placement, metagenome peptides were aligned with the aligned reference sequences using the package PaPaRa (*29*) version 2.5. A maximum likelihood placement was carried out with EPA-ng (*30*), version 0.3.8, with an amino acid substitution model predicted with IQ-TREE. Finally, the jplace file was converted into a newick text file using gappa (*31*) version 0.7.1. Placed sequences with branch lengths longer than the longest distance between all pairs of sequences on the reference tree were removed (<1% of the sequences). Removed sequences were confirmed to correspond to a different function after performing Blastp against the GenBank since the closest sequences were classified with both Pfam or TIGRFAM in different protein families.

### Supplementary Figures

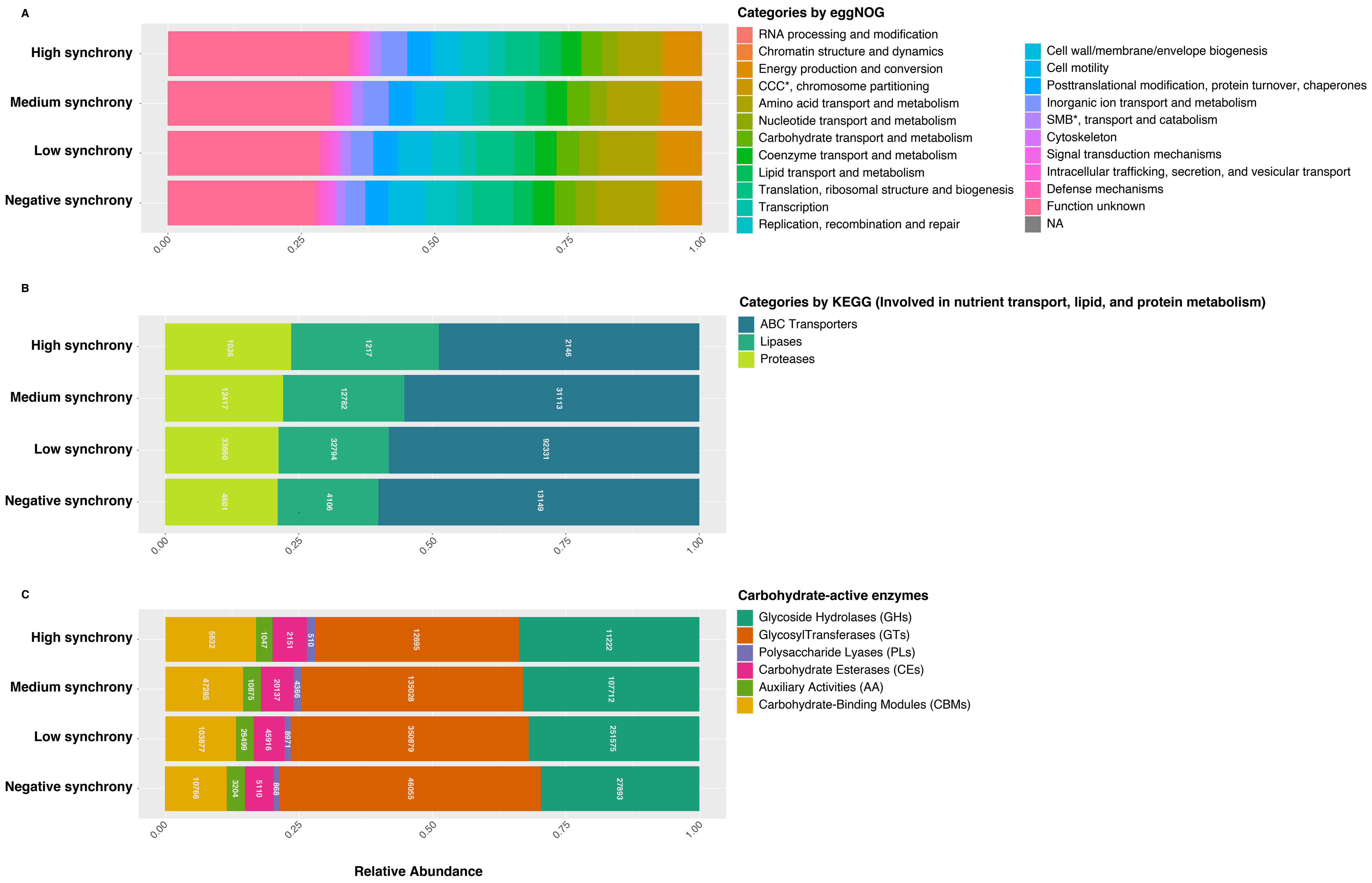

**Figure S1. Similar proportions of genes regardless of synchrony range across all functions.** Functional profiles of BBMO-SOLA genes (ORFs) according to eggNOG, CAZy, and KEGG, analysing 6,375,074 genes with at least 30% of occurrence across all samples. **Panel A.** Relative abundance of genes annotated with eggNOG. SMB, Secondary Metabolites Bio-synthesis; CCC, Cell Cycle Control. **Panel B.** Relative abundance of genes annotated to selected KEGG categories (ABC Transporters, lipases, proteases), since they are crucial for nutrient uptake, protein degradation, and energy production, all of which are fundamental processes in marine biogeochemical cycling. **Panel C.** Relative abundance of genes annotated as Carbohydrate-active enzyme categories.

A

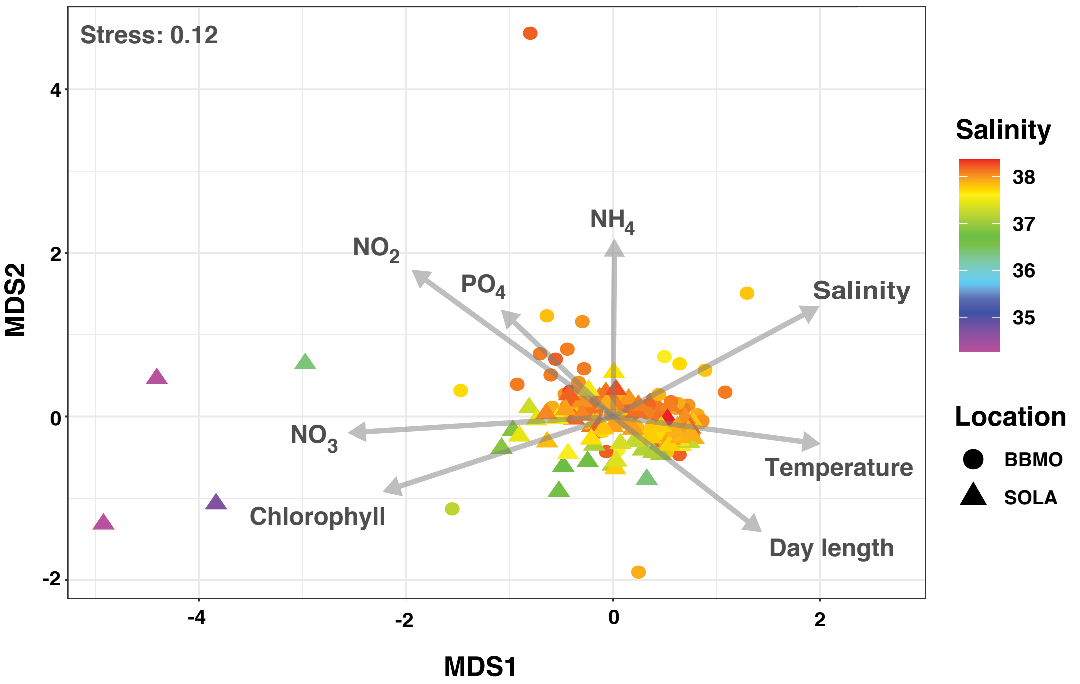


B

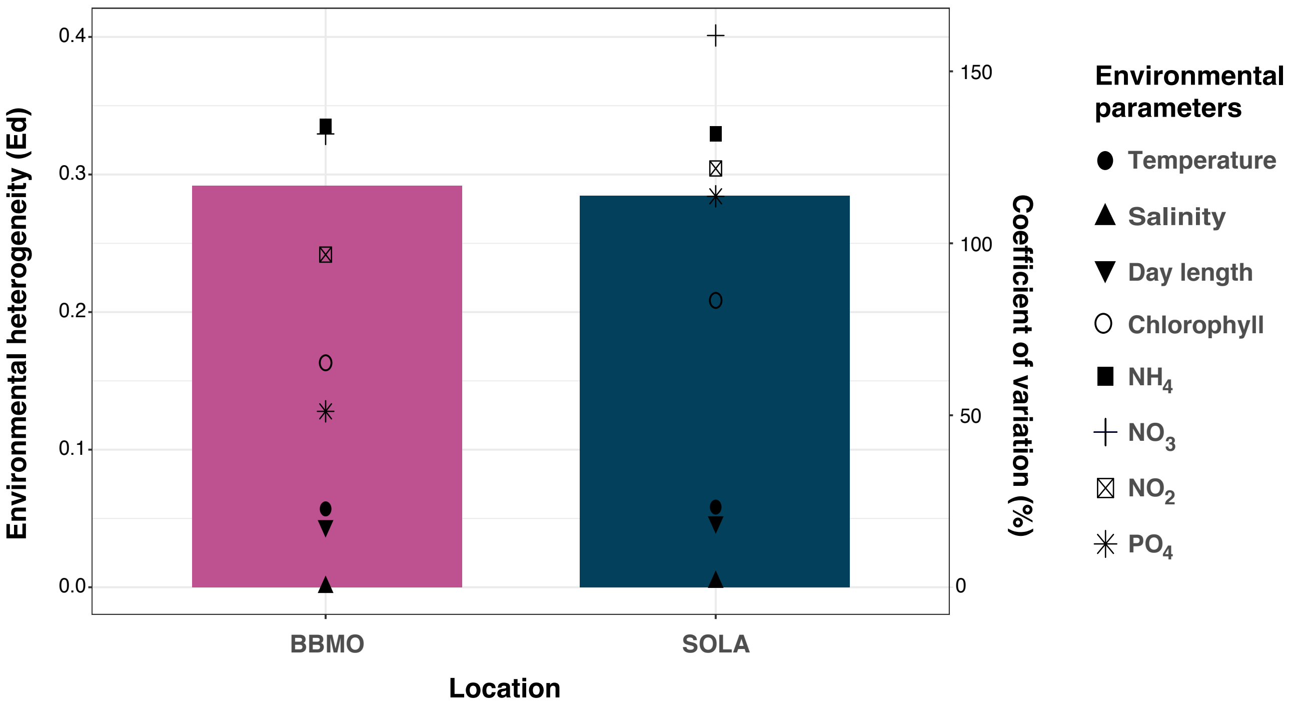

**Figure S2. Similar environmental heterogeneity but different coefficients of variance within BBMO and SOLA. Panel A.** Non-metric multidimensional scaling (NMDS) of the BBMO-SOLA environmental parameters (temperature, salinity, day length, chlorophyll concentration, $NH_4$, $NO_3$, $NO_2$, and $PO_4$). Stress = 0.12. Values were transformed into z-scores before calculating the NMDS. Note that the major differences correspond to lower and higher values of salinity as indicated in a colour gradient. **Panel B.** Environmental heterogeneity in BBMO and SOLA. Environmental heterogeneity is expressed as the average environmental dissimilarity between sites (Ed, coloured bars), and the variability (coefficient of variation, CV%) of the eight parameters is represented with different shapes. Values of the environmental parameters were transformed to z-scores before the analyses.

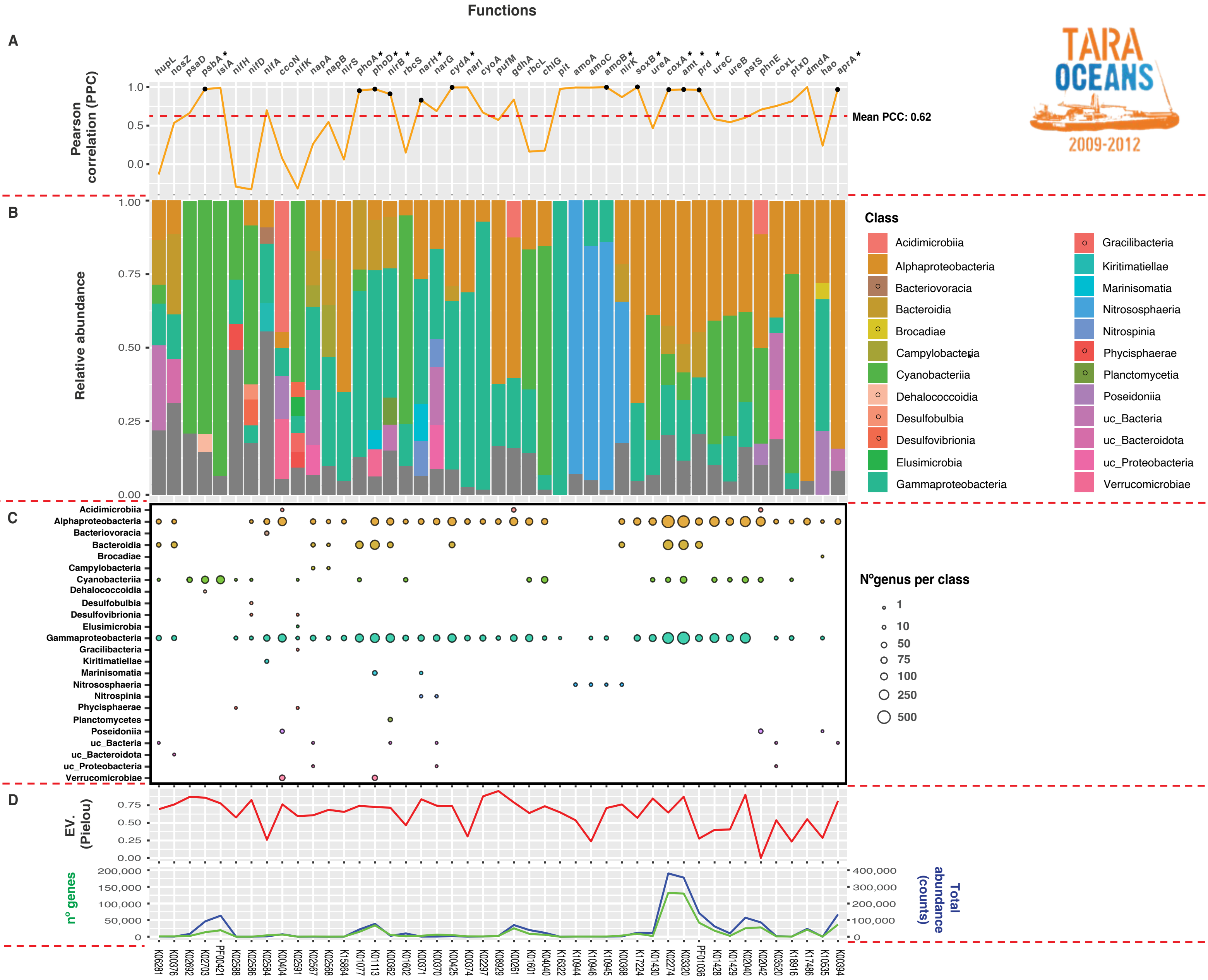

**Figure S3. Taxonomic composition of biogeochemically key functions in the sunlit global ocean (*Tara* Oceans expedition). Panel A.** Pearson correlation coefficient (PPC) between the taxonomic distributions of BBMO-SOLA and the global ocean (*Tara* Oceans) for each function at the Class level as a representation of the similarity in taxonomic composition, indicated with an orange line. Statistically significant correlations are marked with (*). The mean PPC is indicated with a red dotted line. **Panel B.** Taxonomic relative abundance for each function. Taxonomic classes only appearing in the global ocean dataset (*Tara* Oceans) are marked with (o). **Panel C.** Number of genera per class for each function. Colours are the same as in Panel B. **Panel D.** Top. Pielou's evenness index for genes annotated in each function. Bottom. Number of genes (Y axis-left) in green and their total abundance as counts (Y axis-right) in blue for each function.

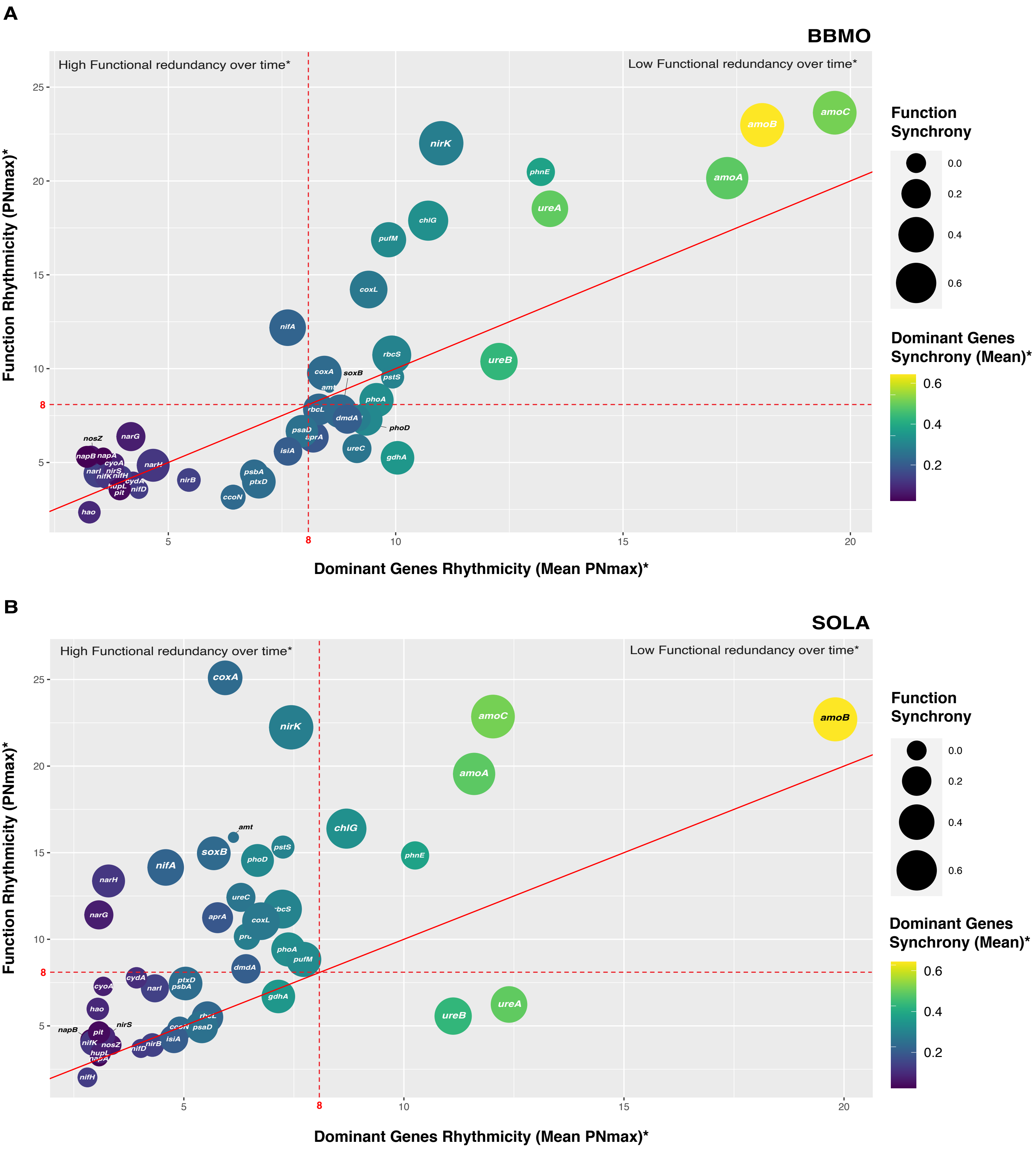
A
BBMO
High Functional redundancy over time*
Low Functional redundancy over time*
Function Rhythmicity (PNmax)*
Dominant Genes Rhythmicity (Mean PNmax)*
Function Synchrony
0.0
0.2
0.4
0.6
Dominant Genes Synchrony (Mean)*
0.6
0.4
0.2
amoC
amoB
amoA
nirK
phnE
ureA
chlG
pufM
coxL
nifA
rbcS
ureB
coxA
soxB
pstS
amt
phoA
rbcL
dmdA
phoD
psaD
aprA
ureC
gdhA
isiA
narG
nosZ
napB
napA
cyoA
narH
narI
nirS
nifK
nifH
cydA
hupL
pit
nifD
nirB
psbA
ptxD
ccoN
hao
B
SOLA
High Functional redundancy over time*
Low Functional redundancy over time*
Function Rhythmicity (PNmax)*
Dominant Genes Rhythmicity (Mean PNmax)*
Function Synchrony
0.0
0.2
0.4
0.6
Dominant Genes Synchrony (Mean)*
0.6
0.4
0.2
coxA
nirK
amoC
amoB
amoA
amt
chlG
pstS
soxB
phoD
phnE
nifA
narH
ureC
rbcS
narG
aprA
coxL
phoA
pufM
dmdA
cydA
cyoA
ptxD
psbA
narI
gdhA
ureB
ureA
hao
rbcL
nirS
napB
pit
ccoN
psaD
nifK
nosZ
isiA
nirB
hupL
napA
nifD
nifH

**Figure S4. Rhythmicity (y-axis) of the main biogeochemically key functions in BBMO (Panel A) and SOLA (Panel B) as a function of the rhythmicity of the dominant genes (x-axis).** The lines at PnMax = 8 indicate the threshold from which a function or gene is considered rhythmic. The bubble size indicates the synchrony of the biogeochemically key functions, while the synchrony of its dominant genes is shown with a colour gradient. Dominant genes are those accounting for >70% of the total abundance of the function. Note that synchrony in both functions and their dominant genes increases with their rhythmicity.

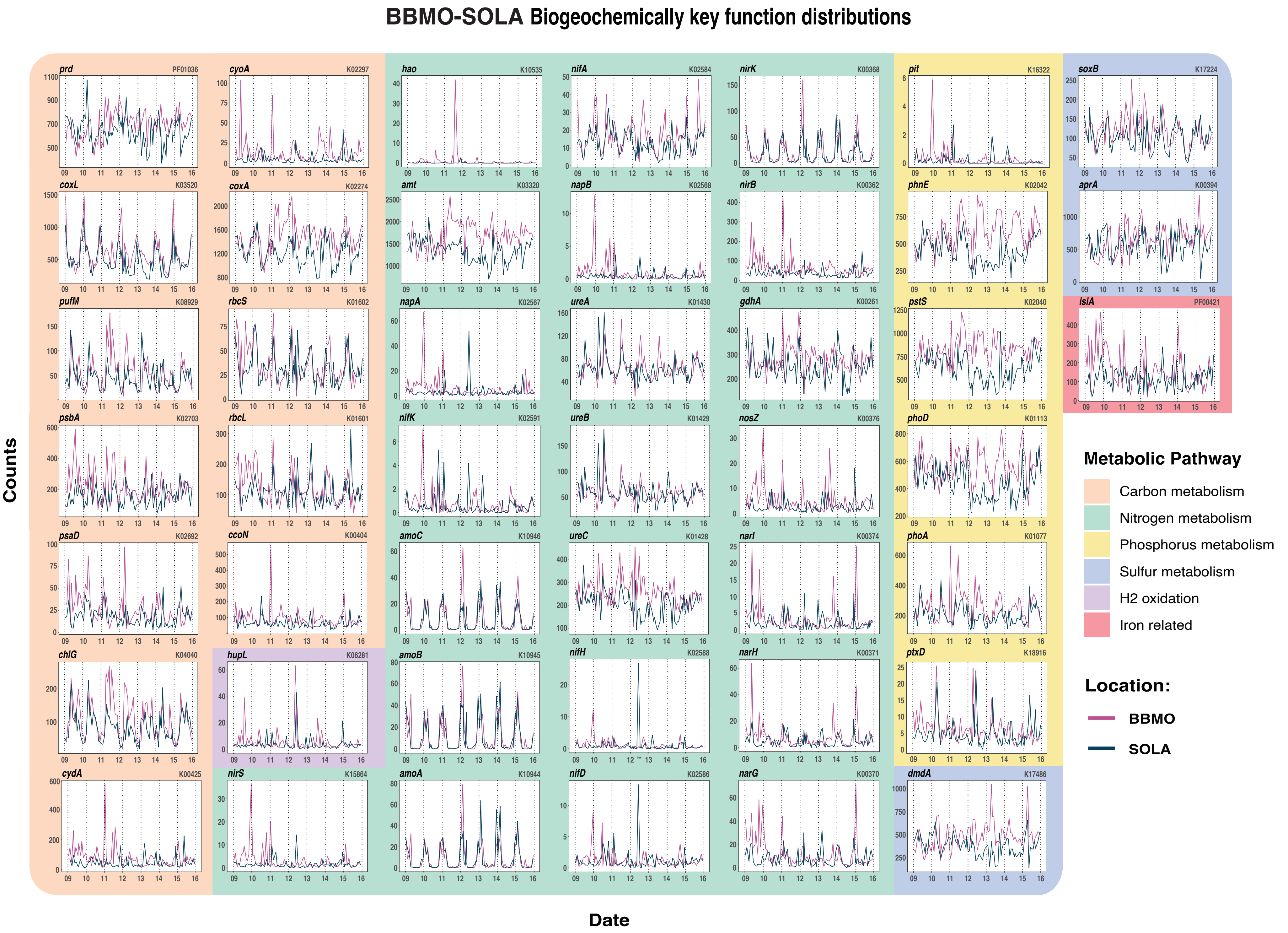

BBMO-SOLA Biogeochemically key function distributions
Counts
Date
prd PF01036
cyoA K02297
hao K10535
nifA K02584
nirK K00368
pit K16322
soxB K17224
coxL K03520
coxA K02274
amt K03320
napB K02568
nirB K00362
phnE K02042
aprA K00394
pufM K08929
rbcS K01602
napA K02567
ureA K01430
gdhA K00261
pstS K02040
isiA PF00421
psbA K02703
rbcL K01601
nifK K02591
ureB K01429
nosZ K00376
phoD K01113
psaD K02692
ccoN K00404
amoC K10946
ureC K01428
narI K00374
phoA K01077
chlG K04040
hupL K06281
amoB K10945
nifH K02588
narH K00371
ptxD K18916
cydA K00425
nirS K15864
amoA K10944
nifD K02586
narG K00370
dmdA K17486
Metabolic Pathway
Carbon metabolism
Nitrogen metabolism
Phosphorus metabolism
Sulfur metabolism
H2 oxidation
Iron related
Location:
BBMO
SOLA

**Figure S5. Abundance of the selected 45 biogeochemical key functions over seven years in BBMO and SOLA.** The metabolic pathway of each gene is indicated with colours. The Y axis shows the sums of the normalized read counts for ORFs belonging to each function as classified with KEGG or Pfam databases. The X axis represents the sampling dates over the seven-year study period.
